# Telomerase RNA gene duplications drive telomeric repeat diversity and evolution in *Andrena* bees

**DOI:** 10.1101/2025.04.29.651168

**Authors:** Christopher Klapproth, Elisa Israel, Abdullah Ahmed, Andreas Remmel, Karl Käther, Franziska Reinhardt, Julian J-L Chen, Sonja Prohaska, Johannes Steidle, Steffen Lemke, Peter F. Stadler, Sven Findeiß

## Abstract

Most organisms in the animal kingdom require a non-coding telomerease RNA (TR) in conjunction with the telomerase reverse transcriptase (TERT) to add telomere tandem repeats to chromosome ends to genomic instability. Recent studies reported an extensive diversity in the sequence of telomeric repeats in some insect species. Our investigation of TR genes in the *Andrena* genus provides convincing evidence for the presence of multiple TR gene copies with different template sequences for synthesis of distinct telomeric repeat sequences in several species. In this study we describe the structure, genomic coordinates and abundance of these TR genes, and correlate our findings with the levels of tandem repeats found in DNAseq data. Based on an analysis of the synthenic context of these newly predicted TR genes, we show evidence for the existence of multiple TR paralogs that diverged during the evolution of the genus *Andrena*. To our knowledge this is the first time such a phenomenon is observed in animals, although recently reported for plants. Interestingly, the comprehensive annotation of all TERT genes found in yet unannotated *Andrena* species shows no corresponding evolutionary changes in related TERT proteins encoded by a single copy gene. Our study suggests an evolutionary mechanism for diversification of telomeric repeat sequences in certain insect species through telomerase RNA gene duplication.

## 1 Introduction

Linear DNA is susceptible to degradation exposed of chromosome ends during the replication process. Telomere structures at the terminal regions serve as a solution to this end-replication problem Levy et al. [1992], Wynford-Thomas and Kipling [1997]. After DNA replication, the addition of characteristic telomeric DNA repeats counterbalances this problem.

Several studies have shown the inter-species diversity of telomeric tandem repeats in eukaryotes Podlevsky and Chen [2016], Závodník et al. [2023]. A substantial diversity of telomeric regions has been described in various organisms, including yeast Cohn et al. [1998], Waldl et al. [2018], Peska et al. [2021], Červenák et al. [2021] and plants Závodník et al. [2023]. This diversity appears to be particularly pronounced in insects Kuznetsova et al. [2020], Zhou et al. [2022], Lukhtanov and Pazhenkova [2023]. However, the reasons behind this atypical variability are currently poorly understood.

The telomerase reverse transcriptase (TERT) plays a fundamental role in the stability of chromosomes in virutally all eukaryotic forms of life Nugent and Lundblad [1998]. With some exceptions, such as drosophilids Pardue et al. [2005], the telomere elongation machinery consists of the TERT protein and a telomerase RNA (TR) that provides the tandem repeat template for reverse transcription Autexier and Lue [2006], Biessmann and Mason [2003], Bryan et al. [2000]. Across eukaryotes, TRs exhibit very poor conservation and differ in form and size Xie et al. [2008], Musgrove et al. [2018]. It follows that TR genes pose a challenge for identification and genomic annotation Podlevsky and Chen [2016]. Furthermore, TR genes in insects have remained elusive until recently Logeswaran et al. [2021], and turned out to be functional although structurally reduced Chou et al. [2025].

Fajkus et al. [2023] identified likely candidates of TR genes in the majority of sequenced Hymenoptera genomes. They used covariance models trained on common structural features in hymenopteran TRs and analyzed potential template regions matching (telomeric) tandem repeats in order to identify new TR genes. Their study suggest a switch of TR transcription by RNA polymerase II towards RNA polymerase III, the biogenesis pathway typically observed in plants Fajkus et al. [2023], Zavodník et al. [2023]. Whether or not there is a connection between large diversity of telomeric regions and a switch in polymerase usage remains to be determined. Reduced TR genes in Lepidoptera were shown to be fully functional Chou et al. [2025]. They share the loss of an H/ACA domain with their hymenopteran counterparts, but are transcribed by RNA polymerase II Chou et al. [2025].

Fajkus et al. [2023] assumed the existence of exactly one TR gene per species and did not investigate the presence of multiple paralogs and corresponding telomeric repeats. There is, however, no particular reason to assume that TR genes exist as single copy genes. For plants, the existence of TR gene paralogs has been reported recently Závodník et al. [2023].

Computational methods dating back as far as the late 90s attempt to identify putative telomeric repeats based on DNAseq data Benson [1999]. The approaches typically rely on statistical analysis of overrepresented tandem repeats in DNAseq data.

In this study, both homology search techniques and analysis of large-scale DNAseq data were used to search for putative TR candidates in species with currently unannotated genomes. For the first time, we present computational evidence for the existence of TR gene paralogs in insects, more precisely in multiple *Andrena* species. Furthermore, 5’-RACE experiments in *Andrena hattorfiana* confirm the existence of at least two independent TR copies. The observed co-existence of multiple TR transcripts, presumably specifying diverse telomeric repeats even within one species, adds another layer of complexity to telomere repeat evolution and the underlying mechanisms.

## 2 Results

### 2.1 Identification of TR genes in *A. dorsata*

Using the telomerase RNA (TR) sequence annotated by Fajkus et al. [2023] and the encoded important features essential during telomere tandem repeat elongation (Figure 1), we identified eight highly similar genetic loci, denoted as AdorTR-1 to AdorTR-8, with Blat in the genome of *A. dorsata*, Figure 2 and Table S1. These loci are spread over the three assembled chromosomes of *A. dorsata* (Figure 2a), and specify a diverse set of five potential template regions resulting in two distinct tandem repeats, Figure 2b. Of the five identified TR template variants, three are presumed to be nonfunctional on the template level due to the lack of functional annealing sites, i.e., AdorTR-2, AdorTR-5 and AdorTR-8.

**Figure 1:**
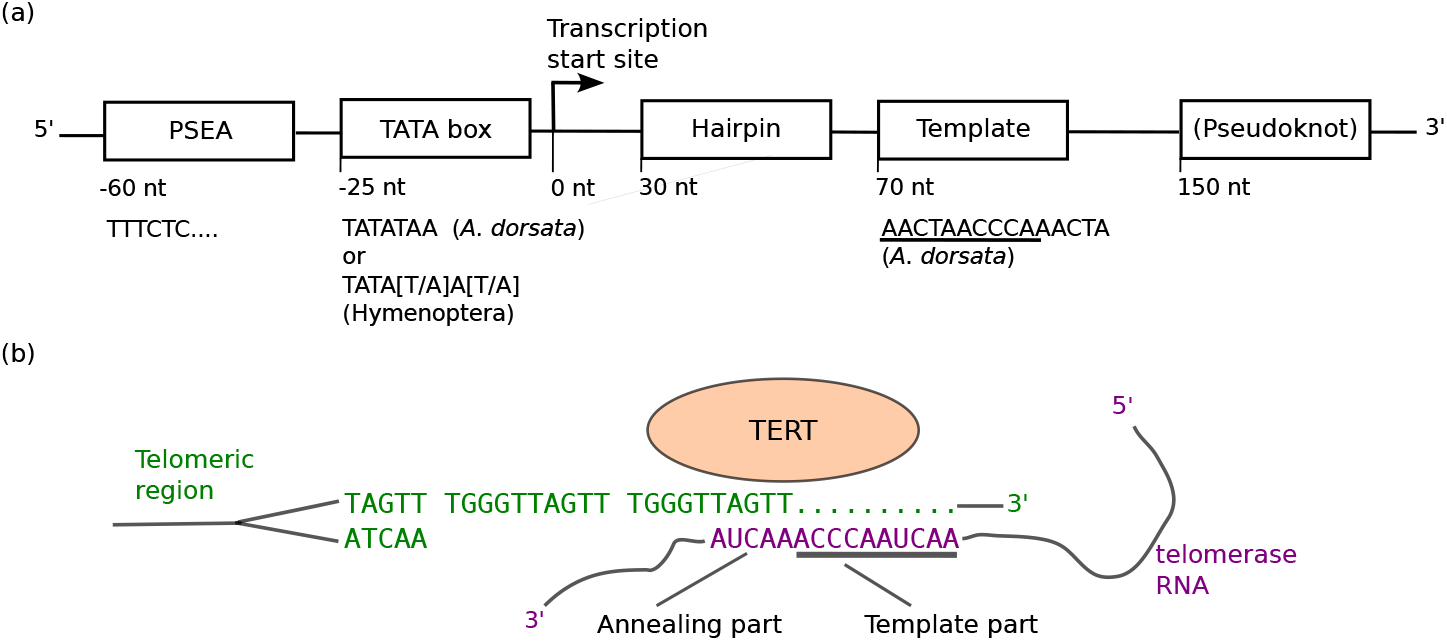
Accepted features of telomerase RNA (TR) genes and their function. (a) Schematic representation of the characteristic structure of TR genes in *A. dorsata* and Hymenoptera in general, according to Fajkus et al. [2023]. Transcription start site is located *∼* 25 nt downstream of highly conserved TATA box and PSEA motifs, indicating an RNA Pol III promoter. The hairpin structure starting around position 30 is highly conserved and found right before the start of the TR template. Fajkus et al. [2023] proposed a pseudoknoted structure at the 3’-end of the TR gene. Sequences reported in *A. dorsata* and Hymenoptera in general are shown below. (b) Illustration of essential components during telomere tandem repeat elongation. The TERT protein complex binds the TR which has the template site for reverse transcription. The template site consists of cyclic permutations of the tandem repeat, forming an annealing and a template part.

**Figure 2:**
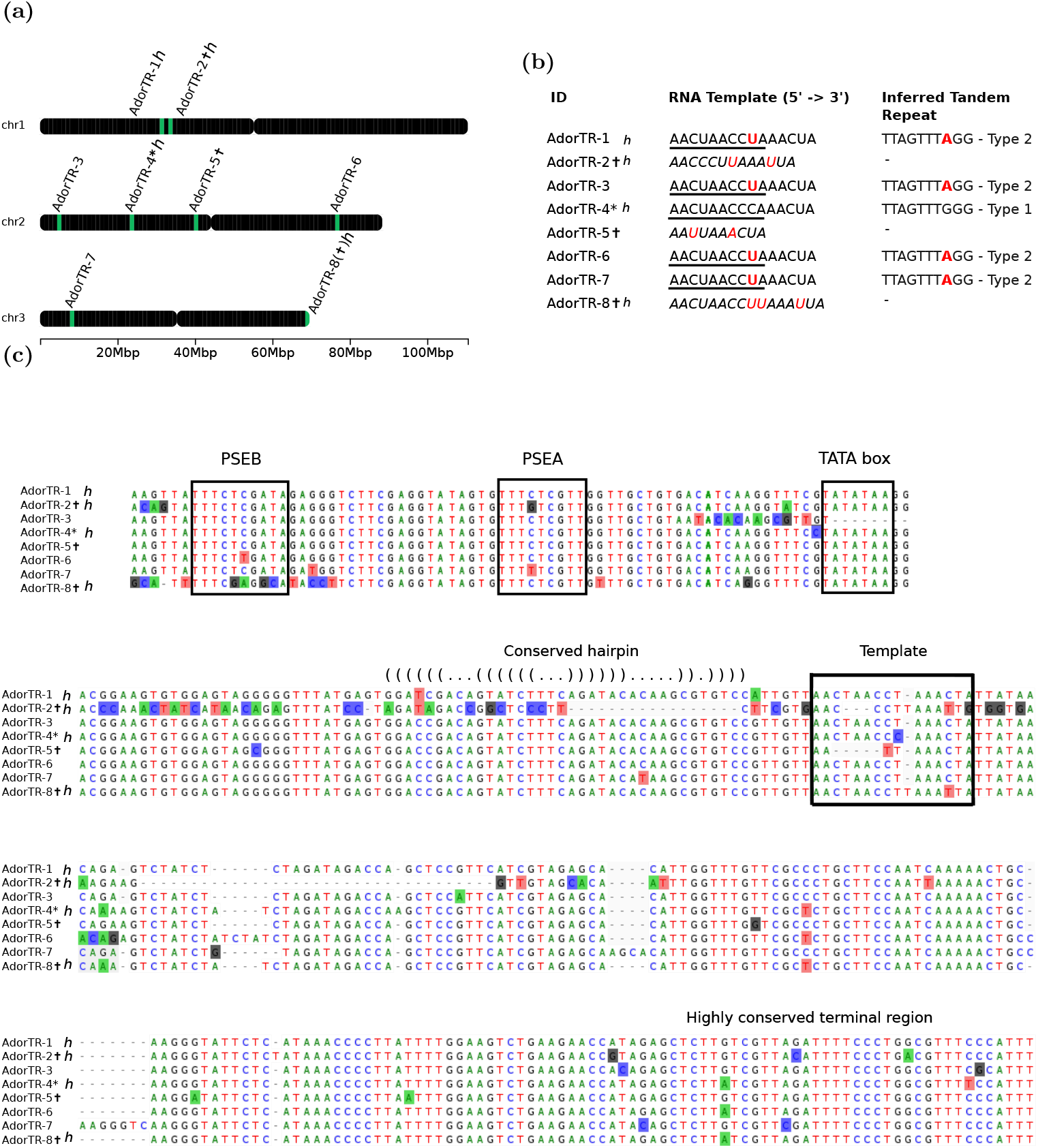
Overview of the TR genes predicted in *A. dorsata* denoted as AdorTR-1 to AdorTR-8. The gene initially annotated by Fajkus et al. [2023] served as the starting point and is therefore marked with an asterisk. (a) TR gene locations along *A. dorsata’s* three assembled chromosomes. (b) Overview of RNA template sites and therefrom inferred tandem repeats. Mutations according to AdorTR-4* are highlighted in red and the template part is underlined, compare Figure 1. AdorTR-2 is notably shorter and misses a conserved hairpin structure otherwise present right before the template region (compare Figure 1). In both AdorTR-2 and AdorTR-5 the template region is partially deleted. As these templates form no viable annealing site for corresponding tandem repeats, the respective genes are presumed to be defective (marked with a t). In AdorTR-8 the template region is modified by the insertion of one additional Uracil nucleotide, potentially breaking the annealing site (compare Figure 3). We furthermore verified our results on an alternate haplotype assembly and identified 4 TR RNA genes corresponding to those annotated in the chromosome level assembly. They are marked with an *h*. (c) Multiple sequence alignment of the eight predicted TR sequences in *A. dorsata*. A high degree of conservation is observed for all features indicated above. A notable property of the predicted TR genes is the highly conserved terminal region, potentially associated with the formation of a pseudoknot.

The inferred tandem repeat TTAGTTTGGG, hereafter also refereed to as Type 1, is the only one recognized by Fajkus et al. [2023] as such, reverse transcribed from the TR template AACUAACCCAAACUA and unique to their predicted gene, AdorTR-4*. The most frequent variation we observe is the template AACUAACCUA present in four TR genes, i.e., AdorTR-1, AdorTR-3, AdorTR-6 and AdorTR-7, and potentially resulting in telomeric repeat TTAGTTTAGG, Type 2. All of these TR genes are otherwise highly conserved on the sequence level, Figure 2c.

Furthermore, a highly conserved predicted RNA Pol III promoter, consisting of a TATA box and at least a proximal sequence element A (PSEA), is present immediately before all sequences except of the AdorTR-2 gene, Figure 2c and Figure S7.

AdorTR-2 is most likely pseudogenic because of its shorter length, a partially degenerate template region with a C*→*U mutation close to the 3’-end and a U insertion, and the missing TATA box motif, Figure 2c. AdorTR-5 is very similar to AdorTR-4* in sequence and structure, compare Figure 2, but misses a five nucleotide long section (six nucleotides when compared to AdorTR-8) in the middle of the TR template. AdorTR-5 and AdorTR-8 also carry point mutations and in case of AdorTR-8 even an insertion within the template region. Additionally, the PSEB of AdorTR-8 is notably mutated. Due to these changes, we assume both to be not functional as templates, similar to the situation found in AdorTR-2.

Taken together, the TR templates of *A. dorsata* are of higher complexity than reported to date in most other animal species, in particular when compared to mammals. This trait is shared with many other Hymenoptera, which also show unusually high TR diversity between species Fajkus et al. [2023]. Interestingly, multiple paralogous TR genes were only recently reported for plants Závodník et al. [2023].

### 2.2 Tandem repeats in DNAseq data of *A. dorsata*

Our hypothesis of concurrent expression of multiple TRs implies that raw DNA sequencing data should contain large numbers of tandem repeats corresponding to the two presumably functional template variants listed in Figure 2b. The two predicted tandem repeats of Type 1 (TTAGTTTGGG)_*n*_ and Type 2 (TTAGTTTAGG)_*n*_ indeed are among the most frequent repetitive sequence patterns, Figure 3. The former corresponds to the template predicted by Fajkus et al. [2023], while the latter matches the template of AdorTR-1, AdorTR-3, AdorTR-6 and AdorTR-7.

**Figure 3:**
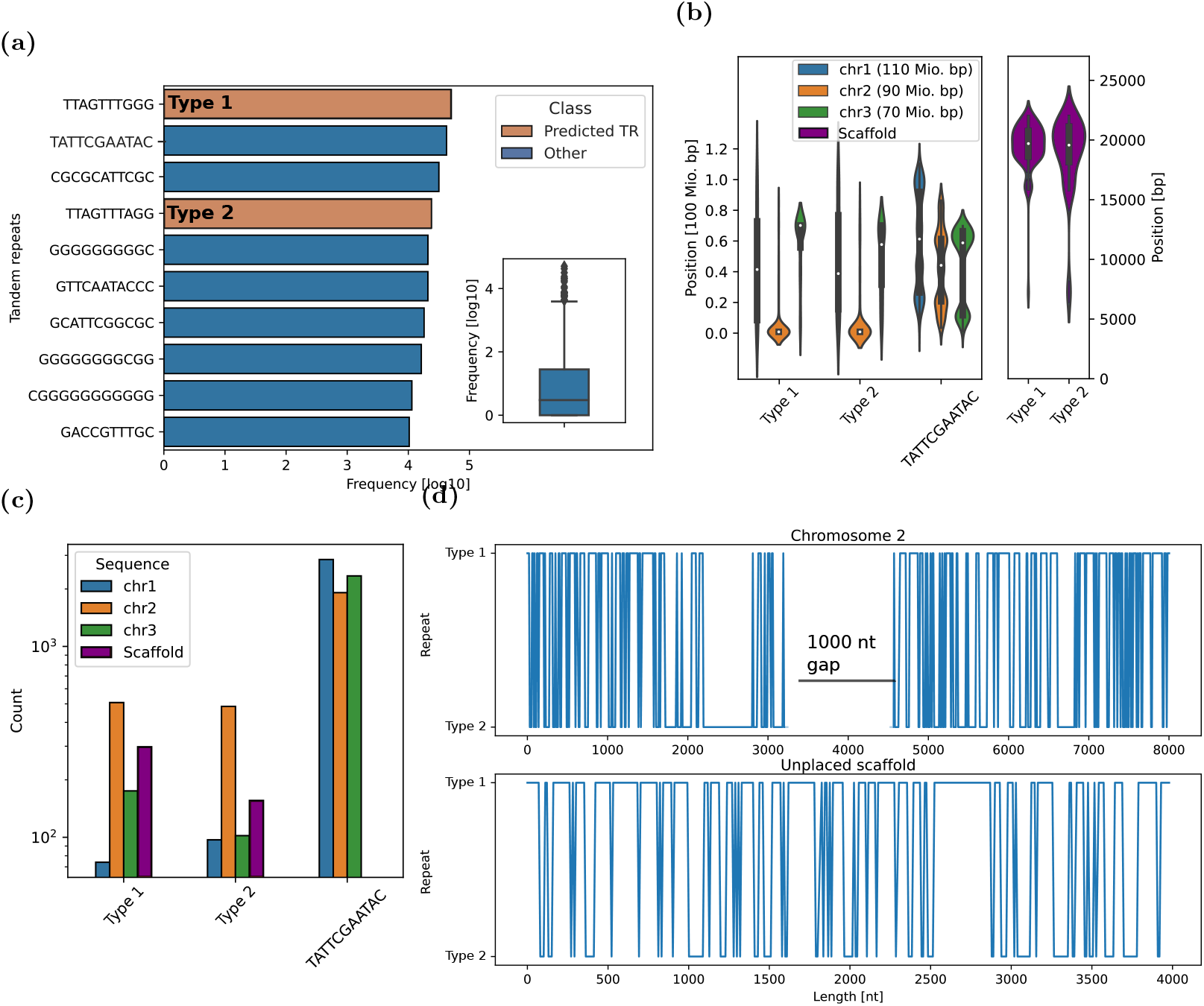
Tandem repeats extracted from raw *A. dorsata* DNAseq (top) and genomic data (bottom). (a) Frequency of the predicted telomeric tandem repeats (TTAGTTTGGG)_*n*_ - Type 1, (TTAGTTTAGG)_*n*_ - Type 2, which are based on the corresponding templates of discovered TR genes is shown in orange. Other highly significant tandem repeats found in the analyzed Hi-C DNAseq data are shown in blue. The box plot in the inset illustrates the frequency distribution of the blue highlighted tandem repeats in general. Most of these repeats are of low complexity. A notable exception is the tandem repeat TATTCGAATAC with unknown function. Repeating the analysis on 10X Genomics Illumina data yields very similar results, compare Figure S1. (b) Distribution and frequency of candidate tandem repeats and the additional complex tandem repeat TATTCGAATAC. Note that the positions obtained here are likely not the complete picture, due to the partially incomplete genome assembly of *A. dorsata*, which currently encompasses many unplaced scaffolds. The largest continuous scaffold was included in the analysis (right side), and yields a notable clustering of two candidate repeats at the 3’-end. (c) Frequency of tandem repeats on the three assembled *A. dorsata* chromosomes and the unplaced scaffold CAKMYH010000194. (d) Alternating pattern of the two tandem repeats found in two longer stretches of presumably telomeric regions on chromosome 2 (5’-end) and the unplaced scaffold (5’-end). The continuous tandem repeat stretch on chromosome 3 is much shorter, but displays basically the same composition, Figure S2.

Comparing these findings with other strongly overrepresented tandem repeats, a lack of sequence complexity in the majority of the remaining repeats is notable, Figure 3a. Most of these repeats are variations of repetitive GC-rich regions. A notable exception is the repeat TATTCGAATAC, which is about as abundant as predicted telomere repeat sequences in the DNAseq data, Figure 3a, and in fact more abundant in the genome assembly, Figure 3c. To analyze the abundance and genomic positioning of this sequence, an exact pattern search against the assembled genome was performed. While TATTCGAATAC seems to be relatively evenly spread along chromosome 1 and 2, there is a notable clustering towards the ends of chromosome 3, Figure 3b. However, no long and continuous regions of this tandem repeat or clustering with other candidate telomeric tandem repeats could be found. It is thus difficult to discern if and how abundantly present this repeat is in telomeric sequences, as the genome assembly lacks the majority of terminal tandem repeat regions of the chromosomes. This fact becomes evident when analyzing the occurrence of the predicted telomeric tandem repeats of Type 1 and Type 2. Both show substantial clustering towards one end of chromosome 2 and 3, respectively.

It is very likely that the presented picture is not complete as *A. dorsata* has more than 100 unplaced genomic scaffolds. Placing high-density tandem repeats correctly in a genome assembly is a rather complicated task. However, at least one scaffold (CAKMYH010000194) shows a high density of tandem repeats, Figure 3c. In the genomic assembly used two stretches of almost uninterrupted repeats can be found, Figure 3d. In particular, we observe roughly 8000 nt at the 3’-end of chromosome 2, with an interruption of around 1000 nt, and 4000 nt on the scaffold without interruption. We analyzed both regions for any repeating patterns and observed that both tandem repeat types, corresponding to TR candidates, seem to alternate randomly, Figure 3d. A final conclusion regarding the actual distribution of the tandem repeats in the *A. dorsata* genome, however, is impossible without experimentally verifying the terminal regions of all chromosomes.

## 2.3 TERT gene annotation in 12 sequenced *Andrena* species

To investigate if the TERT protein and thereby the underlying mechanism of telomere elongation is affected by the observed diversity of TR genes, we annotated the TERT protein coding gene in *A. dorsata* and the eleven other sequenced *Andrena* species, Table S2.

The TERT gene structure, identified utilizing `ExceS-A` Reinhardt and Stadler [2022], consists of 10 coding exons in *A. dorsata* and in fact most analyzed *Andrena* species, Figure 4. This property is also found in other Hymenoptera species, in which the gene is annotated, e.g. *Apis mellifera*. However, variations exist in Hymenoptera, with *Solenopsis invictus* for example having only 7 exons. Observed exceptions in *Andrena* are *A. bicolor* and *A. marginata*, in which no complete match for exon 1 and 2 could be found. In *A. marginata*, we identified a partial match for exon 2, however, the first half is missing or does encode premature stop codons. Instead an alternative start codon exists in the sequence otherwise matching exon 2 or 3, respectively. It is unclear if this is the actual structure of the gene or an artifact of subpar genome assembly. We further observe gene regions encoding functional domains being split into multiple exons, in particular, the RNA binding domain is located on exons 3 and 4. Similarly, the reverse transcriptase domain is usually encoded between exons 6 and 8. The complete essential domain is present in all copies on protein level as identified by `HMMER.`

**Figure 4:**
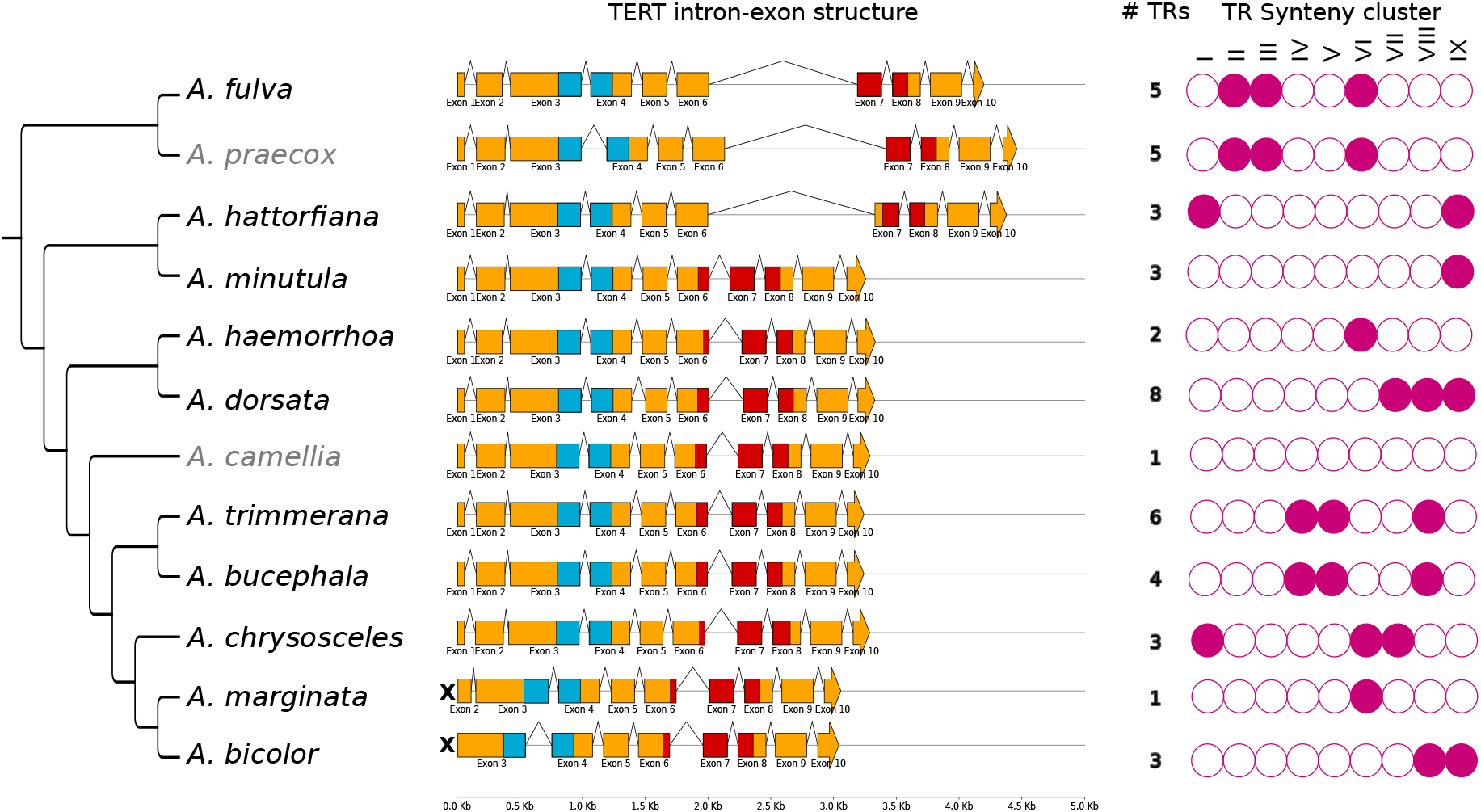
**left**: Phylogeny of *Andrena* species based on the current literature consensus Pisanty et al. [2022], Bossert et al. [2022]. As the positioning of species *A. praecox* and *A. camellia* is currently not known, we estimated their phylogeny based on TERT protein sequence alignments using IQtree Nguyen et al. [2015], Figure S4. Note that the placement of *A. praecox* is much less ambiguous than *A. camellia*. **middle**: Alignment of the predicted TERT gene exon-intronstructure in 12 sequenced *Andrena* species drawn at scale. Analyzing the exon structure, we observe that TERT genes found in *A. bicolor* and *A. marginata* are shortened, denoted with an X in the Figure. Overall, the observed exon structure down to the exon-intron lengths is extremely similar, compare supplementary Figure S3. The functionally crucial RNA-binding and reverse transcriptase domains are highlighted in blue and red, respectively. **right**: Presence or absence of identified telomerase RNA syntenic clusters I-IX, compare the corresponding section. The phylogenetic tree with annotated telomeric repeats can be reviewed in Figure S5. The figure was created using `GenomeViz v.0.4.4` and `iTol` Letunic and Bork [2021].

Furthermore, there is a long conserved intron between exons 6 and 7 found in *A. fulva, A. praecox* and *A. hattorfiana*, lending support to our placement of *A. fulva* in the established phylogeny, Figure 4 and Figure S4.

We verified these results by comparison with an available *A. haemorrhoa* RNA transcript (Genbank: GHFU01005783.1), which corresponds to the TERT gene in this species (verified independently with `HMMER`). Comparing the transcript with the exon sequences found with `ExceS-` shows an almost perfect match with never more than three nucleotides offset per exon boundary and no frame shifts. Furthermore, this comparison reveals the existence of a 79 nt 5’-UTR and a 96 nt 3’-UTR.

In general, these results are in line with those obtained from different haplotype assemblies (data not shown). *A. bicolor* is a special case, as only in the alternate (contig-level) assembly two identical copies of the TERT gene are present. As the corresponding sequences are exactly identical and the observation is isolated to this species, there may be an underlying issue with the contig-level assembly.

## 2.4 Comparison of TRs across the genus *Andrena*

We attempted to reproduce results obtained for *A. dorsata* by searching for TR gene duplicates and differing TR templates in the other eleven sequenced *Andrena* species, see Table S2. In contrast to *A. dorsata*, this process proved more challenging due the high intrinsic variability of insect telomeric tandem repeats even in closely related species Kuznetsova et al. [2020], Lukhtanov and Pazhenkova [2023] and the lack of an experimentally confirmed reference sequence for many of these cases. Fajkus et al. [2023] provide three additional putative TR genes in their supplementary data for the species *A. minutula, A. hattorfiana* and *A. haemorrhoa*. These reported sequences were used alongside the varying putative *A. dorsata* TRs to comprehensively annotate homologs in *Andrena* species.

At least one putative TR with a corresponding genomic tandem repeat was identified for each species, Table 1. In the species for which Fajkus et al. [2023] provided TR gene annotation, we found at least one additional locus. Compared to each other, many of the identified TR candidates show the high diversity typical for insect telomeric tandem repeats within as well as between species. For many analyzed species, evidence for multiple coexisting telomeric tandem repeats could be found, supporting that this phenomenon is not limited to *A. dorsata*. Similar to such cases in *A. dorsata*, there are also a number of putative TRs with no matching tandem repeats found in DNAseq data, indicating that these genes are no longer functional, Table 1. In general, identified telomeric tandem repeats tend to conserve the pattern of starting with a `TT` or `TTA` motif and ending with one or multiple G, even though clear exceptions exist, e.g. *A. fulva* (Table 1), as is also the case in the broader family of insects Lukhtanov and Pazhenkova [2023].

**Table 1:**
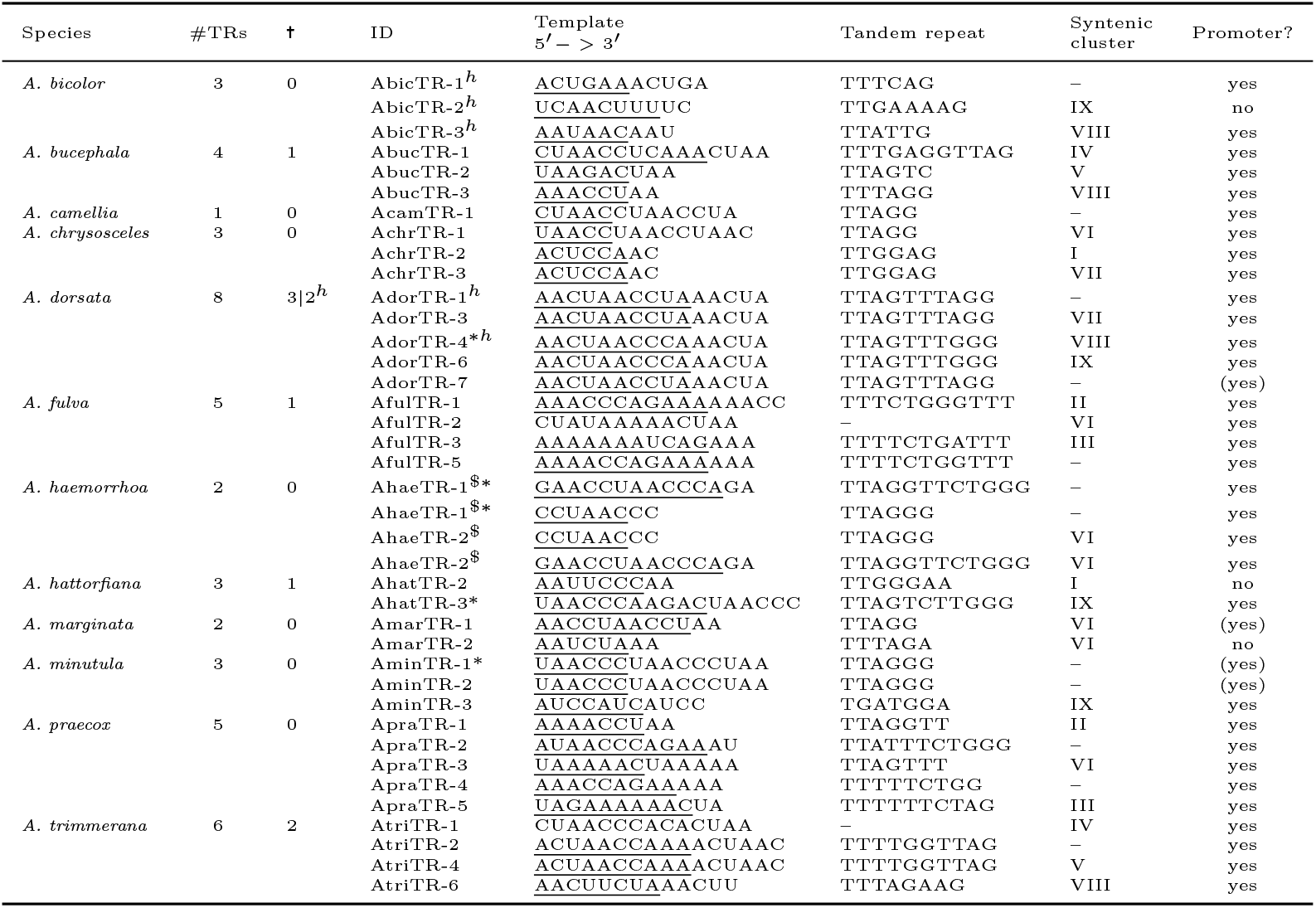
List of putative TR genes, their respective template and inferred telomeric tandem repeats with matching DNAseq evidence. The template part (underlined sequence), compare Figure 1, matches a cyclic permutation of the tandem repeat. A complete list of all potential matches between putative TRs and tandem repeats is available in the supplementary material. Not for all putative TRs such tandem repeats could be identified on DNA level. Due to high divergence of aligned template sites, in a few cases, a faulty alignment may be suspected. However, alignments are usually much better *around* the template region than the template itself. Syntenic information to mitigate this problem was used, denoted as clusters I-IX. Sometimes nearly identical gene copies are found, which do not match any syntenic clusters. We assume them to be recent duplications counted in the copy number column. Putative TRs that do *neither* match any abundant tandem repeats *nor* belong to any identifiable syntenic clusters are omitted, but counted and denoted in the column headed with t. *A. haemorrhoa* is a special case in that the putative template would potentially match two different identified tandem repeats of different lengths at the same time (marked in the table with $). Furthermore, presence and absence of a complete RNA Pol III promoter site, consisting of two proximal sequence elements Hernandez Jr et al. [2007] and a TATA-box (compare Figure S7), is indicated with ‘yes’ and ‘no’, respectively. The presence of a promoter consisting of mutated or even deleted canonical motifs is indicated by ‘(yes)’. Note, that this list is probably non-exhaustive, as our methodology does not account for cases such as the atypical positioning of templates or unusually structured TRs. We highlighted TRs corresponding to seqeuences already annotated by Fajkus et al. with an *. Further, presence of a TR in an available alternate haplotype assembly is indicated (h).

Of the identified matches, a few are of particular note. *A. trimmerana* shows evidence of a template with one U nucleotide inserted, causing a change of the corresponding repeat pattern from `TTTAGAG` to T`TTAGAAG`. Not only have we found point mutations but also evidence of TRs with very different templates within the same species. Examples of this are AAUAACAAU and AUUCUCAUU templates found in *A. bicolor* or ACUCCAAC and UAACCUAACCUAAC in *A. chrysosceles*. Next, reproducibility of these results on alternate haplotype assemblies available for *A. dorsata, A. bicolor* and *A. hattorfiana* was investigated, Table S2. Notably, the available alternate *A. hattorfiana* assembly seems to be incomplete with just 36.5 Mb, compared to 428.5 Mb in the reference assembly and was therefore excluded from the analysis. In both *A. dorsata* and *A. bicolor* we identified genetic loci corresponding to the putative TR genes identified before. In *A. bicolor* all three TR candidates are also found in the alternate haplotype assembly, Table 1. No copies of TRs with identical templates could be found in the alternate

*A. dorsata* assembly, reducing the total number of putative TRs in the contig assembly to four, Figure 2. Interestingly, only two of these, i.e., AdorTR-1 and AdorTR-4 encoding Type-1 and Type-2 telomeric repeats, seem to be functional according to the results presented here, Table 1.

The analysis of closely related Hymenoptera with a chromosome level assembly, see Figure S6, yielded only evidence for the existence of multiple TR genes in *Lasioglossum pauxillium*. However, quality of available genomic data seems to be generally inferior for these species, also reflected by the high number of 17 copies found in *L. pauxillium*. Analogous investigations of the assembled genomes of 30 bumblebees as well as 10 wasps, Table S3, resulted in exactly one TR gene for each species (data not shown).

## 2.5 Promoter analysis of *Andrena* TR RNAs

We searched for sequence motifs 300 nt upstream of all predicted TR genes. Likely candidates for RNA Pol III promoters, consisting of a TATA box and a proximal element A (PSEA), could be identified for almost all candidate genes, Figure S7. Interestingly, there is a highly conserved additional proximal sequence element (PSEB) upstream in *Andrena* species. AdorTR-3, AbicTR-2 and AmarTR-2 display large mutations within the TATA-box, PSE or both regions. Notably, a mutation or deletion in the TATA box seems also indicative of PSE-related mutations. Comparison of the upstream region of TR gene candidates in closely related Hymenoptera, i.e. *Apis mellifera, Bombus terrestris, Macropis europaea, Lasioglossum pauxillium, Osmia bicornis bicornis* and *Colletes gigas*, shows rather little conservation. Although rather distant in this collection, the two *Apidae* might have all three promoter elements upstream of the respective single copy TR genes, Figure S6 and S7.

The structure of RNA Pol III promoters in *Apis mellifera* and some other insects has been described before Hernandez Jr et al. [2007], Mount et al. [2007] and findings presented here are in line with previous analysis of TR genes in Hymenoptera Fajkus et al. [2023]. Additionally, we report the existence of a PSEB element in *Andrena* species, which is still indicative of the overall structure of a type 3 RNA Pol III promoter Sun et al. [2022].

## 2.6 Synteny analysis of TR gene candidates

The above observations give rise to the question when divergence of TRs in insects, or at least *Andrena*, occurred, after or before speciation or even both, with coexisting TR genes being passed on. To address this question, syntenic conservation of the predicted TR gene loci of all twelve *Andrena* was assessed. A difficulty in establishing phylogenic relationships between TRs lies in the rapid mutation of the template region in insects, making the analysis of genomic neighborhood necessary to identify corresponding homologs. A substantial problem here is the lack of annotation data available for *Andrena* thus far. Therefore, the recently published annotation-free approach for synteny analysis by Käther et al. [2023] was utilized. While synteny groups could not be identified for all putative TR genes, results strongly suggest there being multiple pre-speciation divergences of TR genes in *Andrena* species. In total, nine such synteny clusters, denoted I through IX, were identified, Table 1 and Figure 4. Except for *A. camellia*, at least one syntenic cluster could be identified for each species compared to any other and for most multiple clusters where detected, see Table 1. Since *A. camellia* has a sole TR gene with the canonical TTAGG tandem repeat, it is of lesser interest in this study.

For *A. dorsata*, we depict in Figure 5 the syntenic cluster IX. It also serves to illustrate the very high mutability of the *Andrena* TR gene across evolution, with templates even in related genes ranging in diversity from AACUAACCCAAACUA (*A. dorsata*) to the notably shorter AUCCAUCAUCC (*A. minutula*).

**Figure 5:**
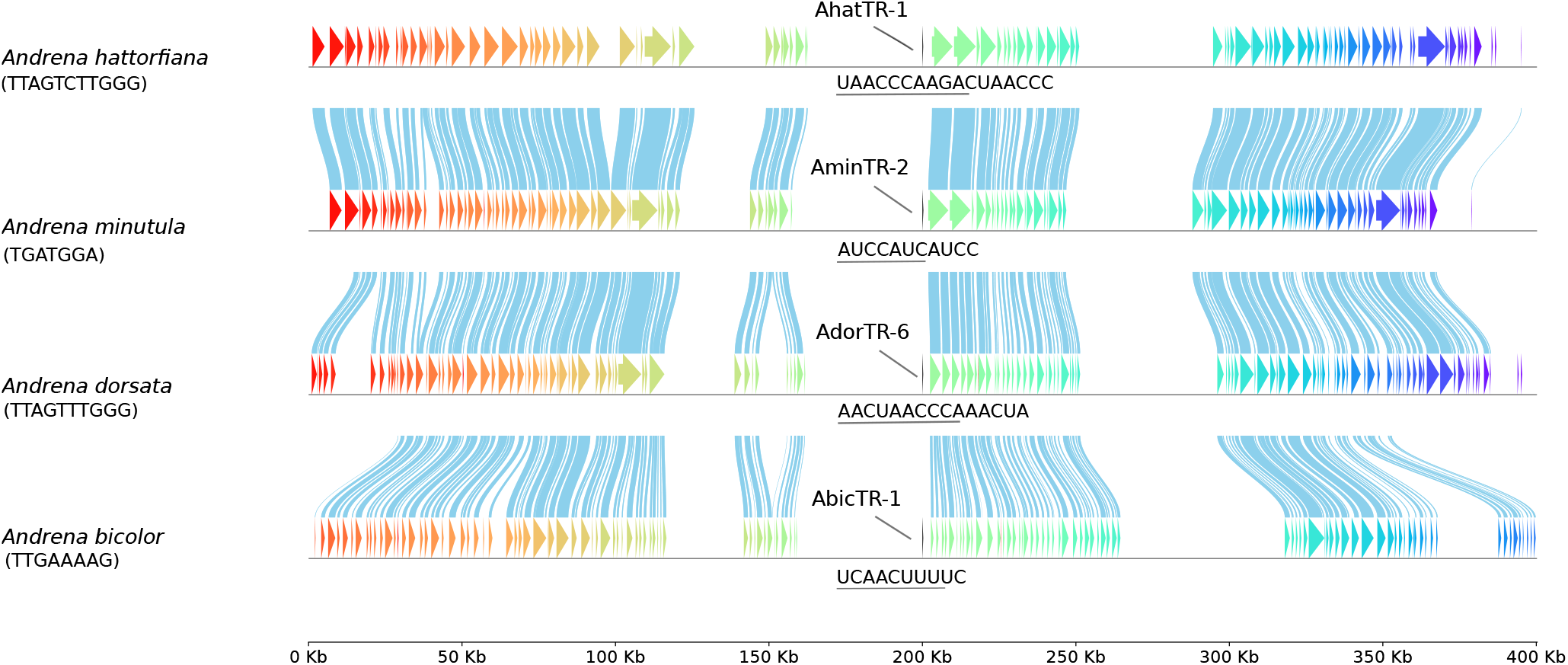
Syntenic cluster IX associated with the telomerase RNA (TR) AdorTR-6 in *A. dorsata* analyzed in a genomic neighborhood of 200 kb in both directions. Pairwise alignments of identified regions of strong similarity are highlighted for better visibility. For better comparison, we show both the TR template (below gene) and the corresponding telomeric tandem repeat identified in the DNAseq data (below species name). Furthermore, genes are identified numerically by their position in Table 1, as well as the associated ID in Table S1 for *A. dorsata*. We observe a drastic change of both the template itself as well as the associated telomeric repeat, illustrating the very strong polymorphism of TRs in *Andrena*. The strong syntenic conservation of this genomic neighborhood in the depicted example, indicates that these four TRs are homologous genes.

To illustrate the existence of multiple coexisting TR genes with individual evolutionary histories, consider the two independent syntenic clusters II and VI, sharing two putative genes each between *A. fulva* and *A. praecox*, see Figure S8. In cluster II, the first gene carries the complex TTTCTTGGGTTT tandem repeat in *A. fulva*, however, presumably due to mutation of the (aligned) template, it specifies the much shorter `TTAGTTT` repeat in *A. praecox*. We observe, however, that the density of repeated `T` nucleotides is still considerable. Notably, the rest of the gene and the synteny of the genomic environment is highly conserved, making an accidental false discovery extremely unlikely. In cluster VI, with a markedly distinct synteny (Figure S8), the TR in *A. fulva* does not carry a tandem repeat (Table 1). However, in this case, the corresponding synteny cluster extends to no less than four additional species including, again, *A. praecox* as well as *A. chrysosceles, A. marginata* and *A. haemorrhoa*. In the latter three, we see the encoding of tandem repeats `TTAGG` and TTAGGG/TTAGGTTCTGGG, respectively. In general, the high mutability of the template region itself emerges as a clear picture, even though the genes and their genomic environment themselves appear to be evolutionarily conserved. To identify potentially conserved ancestral syntenic clusters the above analysis has been extended to closely related Hymenoptera, Figure S6. Despite its evolutionary distance the annotated *Bombus terrestris* TR fits well into cluster IX, Figure S9. In contrast, the evidence for the inclusion of the genes of *Lasioglossum pauxillium* and *Osmia bicornis bicornis* is less pronounced or even scarce in the latter case.

## 2.7 TR gene expression in *Andrena hattorfiana*

To experimentally verify the expression of multiple TR genes in *Andrena* species, we performed 5’-RACE experiments on cDNA templates obtained from *A. hattorfiana*. Our computational analyses indicated that *A. hattorfiana* encodes three TR gene copies; 5’-RACE experiments confirmed the presence of at least two transcripts (AhatTR-1 and AhatTR-3, Figure S10). For both transcripts, the predicted 5’ ends of the two TR candidates are well supported by the RACE data. Notably, AhatTR-3, which specifies the tandem repeat TTAGTCTTGGG, was also annotated by Fajkus et al. [2023] and is likely functional. In contrast, our analysis suggests that AhatTR-1 does not specify a valid template for an abundant tandem repeat, Figure S10. Further experiments are needed to verify TERT binding and to confirm that the telomeric sequences correspond to AhatTR-3. Nonetheless, our findings demonstrate the co-expression of two independent TR gene copies in *A. hattorfiana*.

## 3 Discussion

Genetic loci of high similarity to the TR gene annotated by Fajkus et al. [2023] were identified in *A. dorsata*, which show sequence and strong structure conservation. Based on the findings, we inferred telomeric tandem repeats which would be present in case of transcription and biological activity and are in fact found at the chromosomal ends. We interpret this as strong evidence for the parallel expression of multiple TRs with different templates. Following this, our analysis shows that the situation is not limited to *A. dorsata*, but extends to the majority of species in the *Andrena* genus, with two to three coexisting templates of varying diversity seeming to be the norm. Although recently shown for plants Závodník et al. [2023], to the best of our knowledge, no similar case has been reported for insects and other animals so far. Analogous duplications could not be found in bumblebees or wasps, making these observations unique to *Andrena* so far. It would be of great importance in this context to investigate other related species for TR duplication events, as *Andrena* is among the most speciose genera Pisanty et al. [2022]. Also, the investigation of the evolutionary relationship between the high diversity of telomeric repeats in *Andrena* and the simultaneous development of paralogous TRs remains the subject of future work. As of now, we do not have evidence whether or not this is an issue, but possible heterozygosity of TR alleles might further add to heterogeneity in the respective tandem repeats. Investigating heterozygosity of highly similar multi-copy TR genes (as in *A. dorsata*) will be challenging and might need elaborate experimental setups.

Of particular interest is the strong evidence for the pre-speciation divergence of TR genes obtained from synteny analysis. The concurrent existence of multiple TRs with their own conserved genetic neighborhood and syntenic anchors suggests the coexistence of two evolutionary modes of events: (i) duplication in a common ancestor and (ii) recent duplication of a TR gene after speciation (e.g. AdorTR-3 vs. AdorTR-6 in *A. dorsata*, which notably lack an associated synteny cluster despite high sequence identity).

We hypothesize that there is an evolutionary link between the unusual diversity of telomeric tandem repeats in insects Kuznetsova et al. [2020], Lukhtanov and Pazhenkova [2023], Zhou et al. [2022] compared to many such tandem repeats in vertebrates, yeast and others Meyne et al. [1989], Theimer and Feigon [2006], Kuprys et al. [2013] and the divergence of the otherwise highly conserved TR gene into multiple conserved subgroups. A hypothetical, albeit simple mechanism to explain this abundance, which also matches the data presented here, would be to assume initial duplication events for individual TR genes – note the relative abundance of ‘extra’ copies with no matching syntenic clusters found, see Table 1.

If the conservation signals and the resulting syntnic clusters are taken at face value, there may be one genetic lineage connecting the single-copy *Bombus terrestris* and *Osmia bicornis bicornis* TRs with many of those in *Andrena*. This suggests a rather recent emergence of the multicopy state of *Andrena* TR genes. Given that the multi-copy TR is also present in the halictid *L. pauxillium*, we place the initial duplication event after the divergence of *Apidae* and *Megachilidae* from *Halictidae, Colletidae* and *Andrenidae*. However, the current lack of sequencing data and high quality genome assemblies for closely related species makes this analysis difficult. Furthermore, the presence of only one identified TR in both *Andrena camellia* and *Colletes gigas* suggests that this development is reversible. Nonetheless, determining the exact point of divergence of the multi-copy TRs will be the subject of future research.

Given these extra copies, paired with a high intrinsic mutation rate of the template, this gives evolutionary processes time to experiment with varying templates, of which probably only a fraction become functional telomerase RNA genes. However, assuming a high enough flexibility of the TERT machinery itself, it is quite likely for multiple functional genes to be formed and passed on in the same species. Counteracting this process seems to be a similarly high likelihood of genes becoming defunct through template mutation, effectively turning into pseudogenes – note the presence of seemingly ‘broken’ TRs in *A. dorsata*.

As an additional consideration, we note that a similar observation of multiple coexisting TR paralogs has recently been made in plants Závodník et al. [2023]. Moreover, the predicted RNA polymerase III transcription of TRs in insects is an observation typically made in plants so far, compare Song et al. [2019]. It is remarkable that the existence of both a proximal sequence element A and B seems to be conserved across *Andrena*, Figure S7. Interestingly, PSEB seems to be absent in other Hymenoptera Fajkus et al. [2023]. This indicates a certain mutability not only of the TR sequence, but the entire gene locus including the promoter region, compare the switch from RNA to RNA Pol III described by Fajkus et al. [2021]. Despite our analysis of exemplary species in related Hymenoptera clades, we could only identify multiple TR gene candidates in *Lasioglossum pauxillium*. The presence of 17 putative genes casts doubt on the reliability of the corresponding genome assembly. In general, we observe difficulty with the analysis of related Hymenoptera clades, as two model organisms of *Colletidae* lack either chromosome-level assemblies (*Colletes gigas*) or yield no putative TR candidate at all (*Hylaeus volcanicus*). An in-depth analysis will be the subject of future work. So far, we would not rule out the existence of multiple TRs in closely related Hymenoptera, but it seems rather unlikely in more distant relatives, e.g. Apidae and further.

Considering the question of the function of TR coexistence, we note that the random pattern of switches between different telomeric tandem repeats (Figure 3d) would indicate a constant coupling and decoupling between TERT complex and DNA strand to explain the switch of templates, assuming the TR remains bound to the TERT. This explanation would in turn potentially imply a low TERT processivity.

With our investigation of *Andrena* TERT genes, we can at least hypothesize that the underlying mechanism of telomere elongation itself is highly conserved, as these proteins show no abnormalities in sequence and domain composition compared to known references Nugent and Lundblad [1998], Autexier and Lue [2006]. This was to be expected, as the overall telomerase reverse transcriptase is known to be highly conserved among practically all animals Kanoh and Ishikawa [2003], with very few exceptions, primarily drosophilids Pardue et al. [2005].

Due to the limitations in available data and potentially quality of genome assembly it is impossible to obtain complete certainty of splice sites and transcription boundaries without transcriptome data. Thus, the estimate presented here might differ from the true gene structure. Nevertheless, we are confident that the findings represent a very close approximation of the true sequences and the overall gene locations.

An analysis of available transcriptome data would have improved certainty of our TR analysis. There are, however, three problems: First, to our knowledge, all publicly available RNAseq data sets of *Andrena* use protocols involving polyA-enrichment. In these, the detection of non-coding RNAs with potentially low transcription is difficult if not impossible. Second, due to the high similarity of TR genes within the same species, a distinction by traditional transcript mapping methods will prove challenging. Third, *Andrena* are a very wide spread and abundant species but are, to the best of our knowledge, currently not kept in easily accessible laboratory conditions, compare Pisanty et al. [2022].

To investigate the underlying evolutionary scenario leading to the emergence of multiple TR copies, the essential next step would be to extend the analysis to closely related families. In this case, research would be necessary to pinpoint a species which is more capable of being held in a wet lab.

*Andrena* is an additional group that showcases the diversity of the telomere elongation machinery in eukaryotes that warrants further research to explore underlying mechanisms and implications for biological function. We suggest, therefore, that an experimental verification of the computational results reported here is a worth-while endeavor and the methodology outlined here be applied to other prospective species.

## 4 Materials and Methods

### 4.1 Identification of TR genes in *A. dorsata*

Using the telomerase RNA (TR) gene annotated by Fajkus et al. [2023], we performed `BLAT` Kent [2002] searches on the *A. dorsata* reference genome (Genbank accession GCA 929108735.1, Lin et al. [2023]). Candidate hits were filtered based on their similarity to the query sequence and the presence of a conserved template region from which a telomere repeat sequence could be transcribed. All in-house scripts used in this study are available at github ^1^.

### 4.2 Tandem repeats in DNAseq data of *A. dorsata*

To confirm the presence of predicted telomerase repeats corresponding to TR templates, we analyzed raw DNAseq data (Supplemental Table S2) from *A. dorsata*, screening for tandem repeats. The library was assembled from paired-end reads and originally sampled using a Illumina NovaSeq 6000. Tandem repeats were identified using the program tandem-repeat-finder Benson [1999]. Results were then clustered with the publicly available software `tandem-repeats-merger`^2^, yielding an initial list of candidate repeats. In a second post-processing step, we merged identical candidates that only differed in starting positions and frames. This post-processing was done using in-house scripts made available at github ^3^.

### 4.3 TERT gene annotation in 12 sequenced *Andrena* species

The amino acid sequence of the annotated TERT-like protein (XP.015428996.1) of *Dufourea novaeangliae* was obtained from the NCBI database. This record was derived from the genomic sequence using gene prediction method Gnomon which is part of the NCBI’s annotation pipeline. Compared to *Andrena* it is the most closely related species with an annotated TERT protein in `NCBI. ExeS-A` Reinhardt and Stadler [2022], a splice-site aware split aligner, was applied to annotate TERT genes in all 12 *Andrena* species with sequenced genomes and a chromosome-level assembly, Table S2. First, the amino acid sequence of the *D. novaeangliae* TERT protein was searched against its own genome in order to calibrate the `ExeS-A` parameters. Second, this amino acid sequence was used to search in *A. fulva*, one of the early branching *Andrena* species. The retrieved full-length hit of *A. fulva* was subsequently used as query against the remaining eleven *Andrena* species. For this analysis, the minimal `ExeS-A` score of splice sites was adapted to -14. Otherwise, the program was applied with standard parameters as available on `github` ^4^.

Exon sequences obtained for *A. haemorrhoa* were compared with the RNA transcript under genbank accession GHFU01005783.1, which represents a spliced transcript of the TERT gene in *A. haemorrhoa*. Alignments were created with `BLAT` and exon boundaries compared manually.

The presence of the required reverse transcriptase domain and the RNA-binding domain in all detected protein sequences was validated utilizing `HMMER` Eddy [2011] with HMM-models of the respective domains as obtained from the InterPro database Paysan-Lafosse et al. [2023]. This, however, only verifies the presence of the minimal required TERT protein domains for biological function. Some sources state that the loss of the telomerase essential N-terminal (TEN) domain interferes with telomerase activity in insects, compare Mason et al. [2011]. However, we point out that the TERT genes observed here are highly conserved compared with TERTs annotated in other Hymenoptera clades, which do seem to have telomerase-like activity Fajkus et al. [2023].

### 4.4 TR annotation in other sequenced *Andrena* species

Fajkus et al. [2023] provide three additional putative TR genes in their supplementary data for the species *A. minutula, A. hattorfiana* and *A. haemorrhoa*. These reported sequences were used alongside the varying putative *A. dorsata* TRs as BLAT and blastn search queries against the genomes of *Andrena* species, Table S2. For comparison, we also repeated this analysis for alternate haplotype assemblies of *A. dorsata* and *A. bicolor*. Additionally, we performed a `blastn` search of all candidate TRs in a species against the genome they were identified in, to catch slightly altered copies within the same genome.

Statistically overrepresented tandem repeats based on available DNAseq data were obtained in the same way as for *A. dorsata*. The utilized DNAseq data sets for each species are listed in Table S2. Hi-C Illumina read data was used when available, which was the case for every species except *A. camellia* and *A. fulva*. We presume the available data for those species is of lower quality. Therefore, we cannot guarantee the same level of specificity for these species. Published telomeric tandem repeats for *A. minutula, A. hattorfiana, A. haemorrhoa* and *A. dorsata* were utilized as control for the tandem repeat identification process Lukhtanov and Pazhenkova [2023].

Likely matches between putative TRs and predicted telomeric repeats were identified as follows: For each individual non-redundant tandem repeat we checked if a cyclic permutation of at least length *L* = *l*_*r*_ + 2 could be found in any putative TR, where *l*_*r*_ is the length of one repeat. If so, the RNA is aligned with (a) all other putative telomerase RNAs found in this species and (b) pairwise with the full AdorTR-4* sequence, identified by Fajkus et al. [2023]. The latter alignment was used to identify likely coordinates of the TR template while the former was used to ensure that putative TR templates of the same species would be sufficiently similar. Ultimately this was done to ensure that the TR template was placed in a sensible position relative to other features of the gene. This way, likely false positives of tandem repeats matching at the terminal ends of the RNA could be eliminated. In unclear cases, we assumed that no valid template would be located further downstream than 150 nt within the RNA, as suggested by analysis of the annotated TRs done by Fajkus et al. [2023]. Remaining matches between terminal repeats and putative TRs are listed in Table 1.

### 4.5 Promoter analysis of *Andrena* TRs

To analyze putative promoter regions, 300 nt upstream genomic regions of all predicted *Andrena* and closely related Hymenoptera TRs were obtained and aligned with clustalw2. Motifs were predicted *de novo* using the MEME suite Bailey et al. [2009, 2015] with standard parameters `(-dna -mod zoops -nmotifs 3 -minw 6 -maxw 25 -objfun classic -revcomp -markov order 0)` and compared with established PSEA and TATA-box motifs for RNA Pol III promoters in insects and Hymenoptera Hernandez Jr et al. [2007], Mount et al. [2007].

### 4.6 Synteny analysis of TR gene candidates

Syntenic conservation of the loci was assessed using pairwise local alignments which were further clustered into multiple alignments. The annotation-free approach by Käther et al. [2023] calculates pairwise alignments in a two-step process. First, k-mer counts are used as a proxy to select promising candidate sequences for alignments in each genome. These candidates are then aligned to the most similar sequence in their own genome resulting in a candidate set per genome. Next, these candidate sets are compared between genomes and, upon fulfilling the condition that an alignment between genomes is significantly better than within genomes, pairwise matches are reported. Thus, ambiguity introduced by aligning potentially paralogs or otherwise repetitive loci is avoided. In order to obtain multiple alignments, pairwise ones are extended transitively if they fulfill a locality condition. Candidate sequences are considered nodes in a graph and pairwise alignments as edges Käther et al. [2023]. Each pairwise alignment is assigned a cluster of multiple pairwise alignments which form a connected component of the graph. If there are multiple candidate sequences of one species in a cluster with a gap of more than 5000 nucleotides, the cluster is discarded.

These multiple alignments are then used to cluster the genetic loci themselves by considering the potential multiple alignments in their genomic neighborhood. Hence, all loci which share any syntenic alignment are clustered. This syntenic clustering yields different numbers and types of clusters, depending on the size of the genomic neighborhood considered. Given that the clusters do not change taking a neighborhood of *±* 50 *kb* and *±* 100 *kb* and that the elements in the clusters identified do not change but two additional clusters arise for *±* 200 *kb*, it can be assumed that the clustering is reasonably reliable for *Andrena*. When increasing the genomic neighborhood sizes to *±* 500 *kb* and 1 *Mb*, some loci also change their cluster membership, but this does not change the overall picture leading to our conclusions, see online supplemental.

### 4.7 5’-RACE in *A. hattorfiana*

5’ and 3’ rapid amplification of cDNA ends (RACE) reactions were performed using SMARTer RACE kit (Takara Bio), starting with total RNA obtained from a single specimen that was collected during the summer of 2024 in the Hohenheim area (Germany); the frozen specimen was homogenized in TRIzol (Invitrogen), followed by RNA extraction according to the manufacturer’s protocol. 5’-RACE reactions were performed as nested PCRs, either with degenerate primers with consensus binding to all candidate genes (5’-GRRKTTWAYGAGAATACCCTTSC, 5’-TTTTTDATTGGAAGAAGGGCGA), or with gene specific primers for each of the candidate TR transcripts, Table S4 (AhatTR-1: 5’-AAATGGGAAACGCCAGGGAAAAA, 5’-GCAGTTTTTGATTGGAAGAAGGGCG; AhatTR-2: 5’-AAATGGGAAACGCCAGGGAAAAA, 5’-AGACTTCCAAAAAGAAAGGGGGAGT; AhatTR-3: 5’-ACGCCAATTCTTCACAGGACTTCCA, 5’-ACGAGAATACCCTTCCCAGACTTAGGT).

## 5 Data Availability

Essential in-house scripts used in this study as well as new raw data sets underlying Figures and analysis can be downloaded from: https://github.com/chrisBioInf/TR-RNA-gene-duplication-supplementary-scripts. External data sources, such as SRA accessions, are listed in the supplementary document.

## 6 Acknowledgments

This work was partially funded by the DFG (German Research Foundation) in GEvol SPP (Project Numbers: 502862570 (STA850/60-1) and 503349225 (PR1288/9-1)) and the CRC-1423 (Project number: 421152132), by the Federal Ministry of Education and Research of Germany and by the Sächsische Staatsministerium für Wissenschaft, Kultur und Tourismus in the program Center of Excellence for AI-research “Center for Scalable Data Analytics and Artificial Intelligence Dresden/Leipzig”, project identification number: ScaDS.AI.

## Supplemental Information for

**Table S1:**
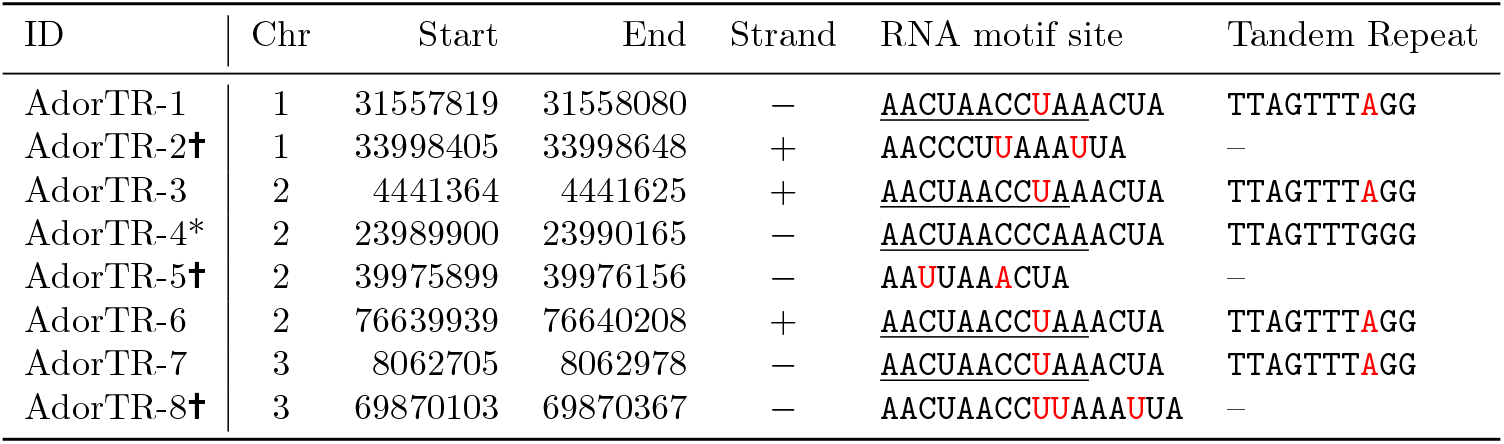
A list of predicted TR genes in *A. dorsata* and their respective genomic locations. The gene AdorTR-4, marked with an asterisk, was initially annotated by Fajkus et al. [2023] and served as the starting point. Motif sites in each predicted RNA that contain the template region from which the telomere repeats are transcribed and the inferred telomere repeat sequence that would be expected are listed. Differences to the sequence predicted by Fajkus et al. [2023] are highlighted in red. Based on the motif sites of gene AdorTR-5 and AdorTR-2 we expect them to be not functional (indicated by t), as both lack a working annealing site due to a C *→* U mutation at position 3 and 11, respectively.

**Table S2:** List of all used *Andrena* species, genome accession, DNAseq data IDs and genome assembly statistics, i.e. genome size, number of chromosomes, number of scaffold, submitter, year and assembly level. For three species, we also investigated alternate haplotype contig assemblies for comparison. WSI = WELLCOME SANGER INSTITUTE; CNU = Chongqing Normal University; ahCon = alternate haplotype contig; chrom = chromosome level assembly; TC = Targeted-Capture; WGS = Whole Genome Shotgun Sequencing.

| Name | Genbank ID | DNaseq | Assembly Statistics | Strategy |
| --- | --- | --- | --- | --- |
| *A. dorsata* | GCA_929108735.1 | ERR7220497 | 277.3 Mb, 3/157, WSI, 2022, chrom | Hi-C |
|  | GCA_929114765.1 |  | 215.9 Mb, 3/157, WSI, 2022, ahCon | Hi-C |
| *A. bicolor* | GCA_960531205.1 | ERR10684073 | 351.7 Mb, 5/325, WSI, 2023, chrom | Hi-C |
|  | GCA_960531515.1 |  | 324.8 Mb, 5/325, WSI, 2023, ahCon | Hi-C |
| *A. bucephala* | GCA_947577245.1 | ERR10123710 | 379.8 Mb, 5/13, WSI, 2022, chrom | Hi-C |
| *A. camellia* | GCA_029448645.1 | SRR23869504 | 340.7 Mb, 7/50, CNU, 2023, chrom | WGS |
| *A. chrysosceles* | GCA_963855975.1 | ERR10818314 | 520.2 Mb, 4/813, WSI, 2023, chrom | Hi-C |
| *A. fulva* | GCA_946251845.1 | SRR13987936 | 461.7 Mb, 7/317, WSI, 2022, chrom | TC |
| *A. haemorrhoa* | GCA_910592295.1 | ERR6054880 | 330.7 Mb, 7/402, WSI, 2021, chrom | Hi-C |
| *A. hattorfiana* | GCA_944738655.2 | ERR9580473 | 428.5 Mb, 7/125, WSI, 2022, chrom | Hi-C |
|  | GCA_944738785.2 |  | 36.5 Mb, 7/125, WSI, 2022, ahCon | Hi-C |
| *A. marginata* | GCA_963932335.1 | ERR11182527 | 373.6 Mb, 6/522, WSI, 2024, chrom | Hi-C |
| *A. minutula* | GCA_929113495.1 | ERR7113537 | 380.4 Mb, 7/358, WSI, 2022, chrom | Hi-C |
| *A. praecox* | GCA_963932275.1 | ERR10501025 | 563.1 Mb, 7/664, WSI, 2024, chrom | Hi-C |
| *A. trimmerana* | GCA_951215215.1 | ERR10395991 | 399.1 Mb, 5/225, WSI, 2023, chrom | Hi-C |

**Table S3:**
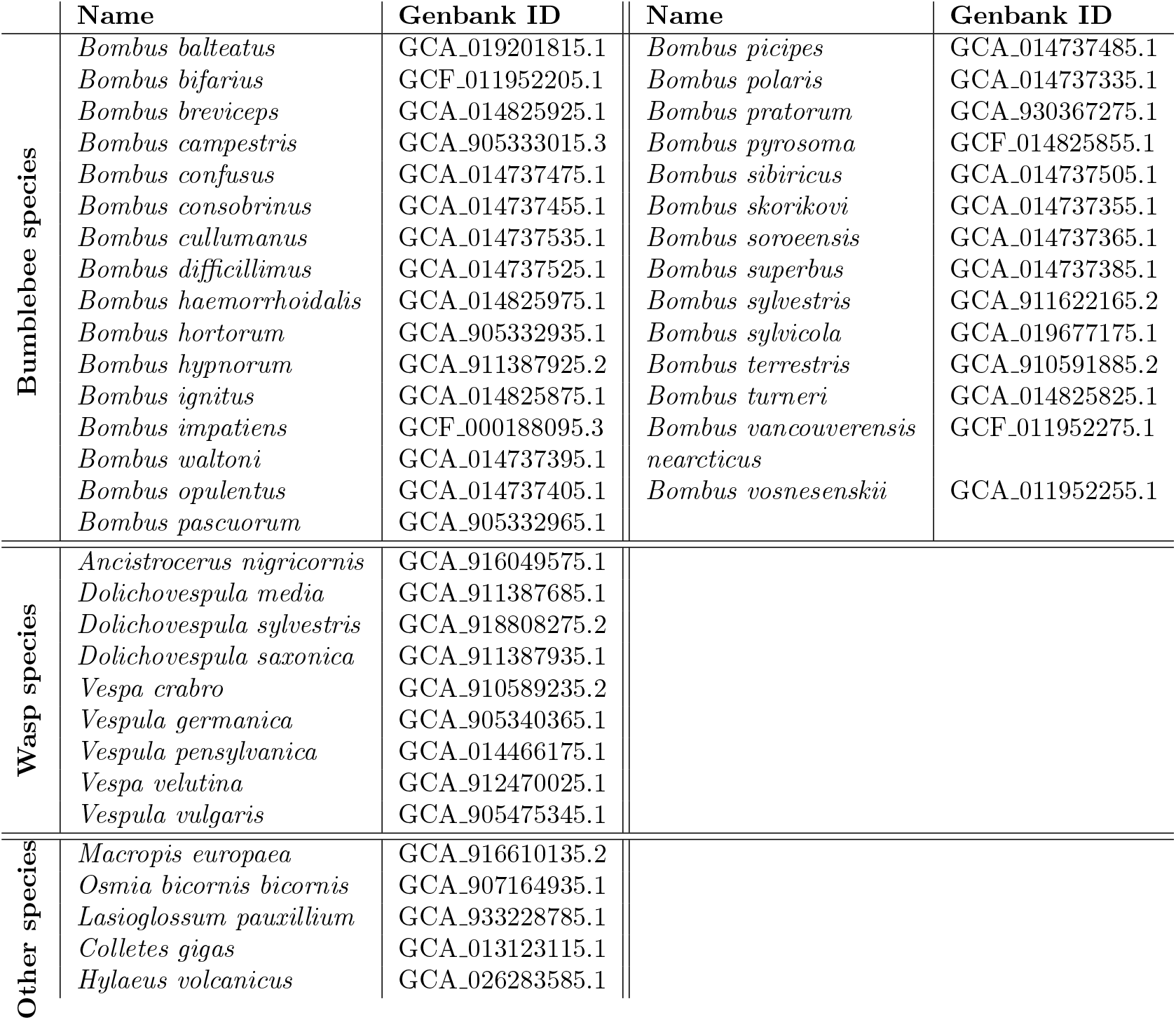
List of bumblebee, wasp species and additional closely related species used in a comparative analysis.

**Table S4:**
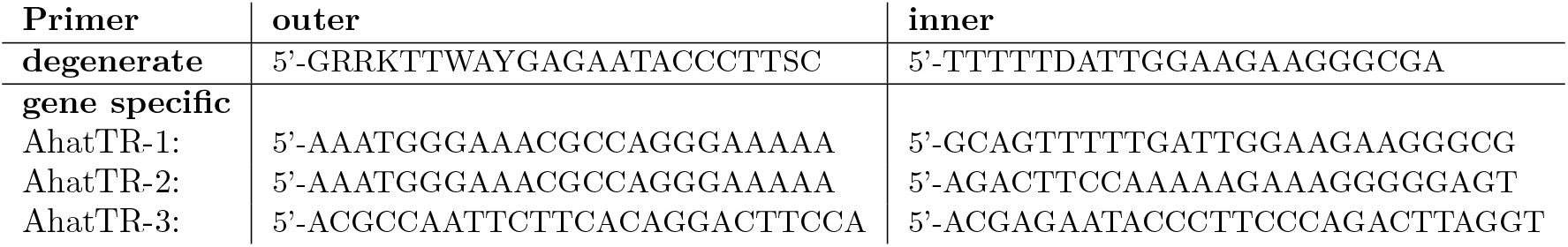
Nested PCR primers applied for 5’-RACE reactions. Either degenerate primers with consensus binding to all candidate genes or gene specific primers for each of the individual candidate TR transcripts were utilized.

**Figure S1:**
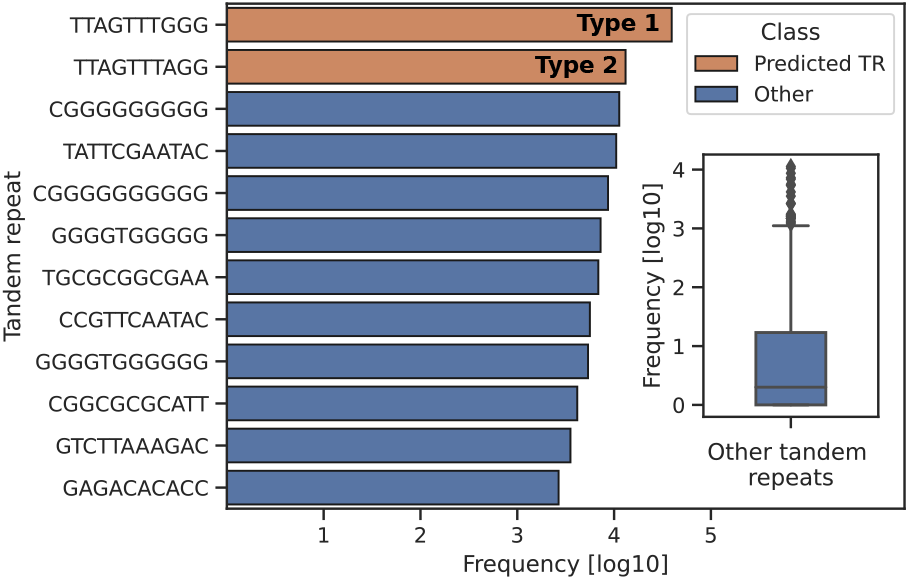
Overrepresented tandem repeats extracted from raw *A. dorsata* 10X Illumnia DNAseq data (ERR7220493-ERR7220496). Color-encoding same as in Figure 3a of the main-text.

**Figure S2:**
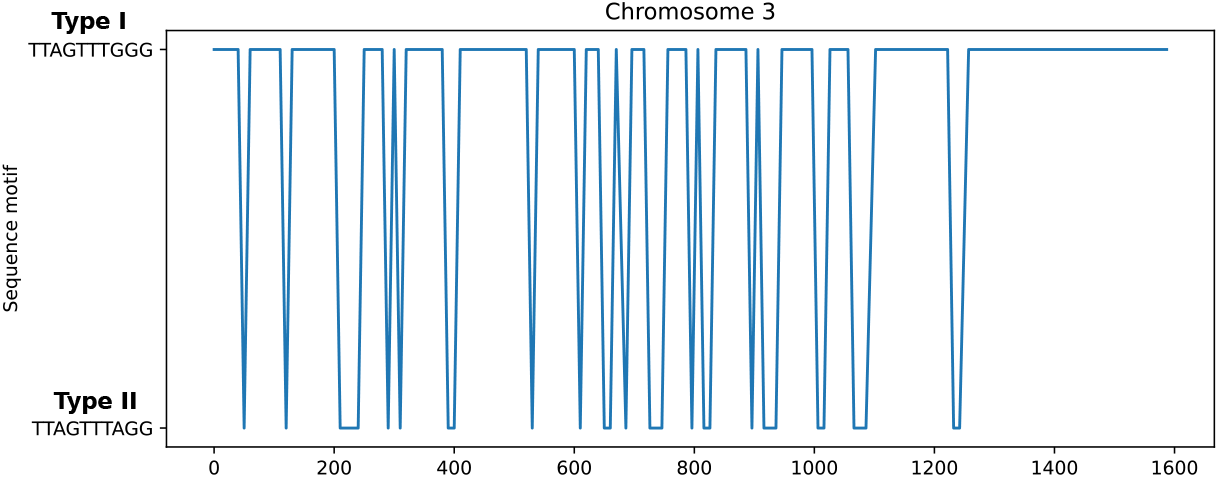
Interchanging pattern of two telomeric tandem repeats at the 3’-end of chromosome 3 in the *A. dorsata* genome assembly.

**Figure S3:**
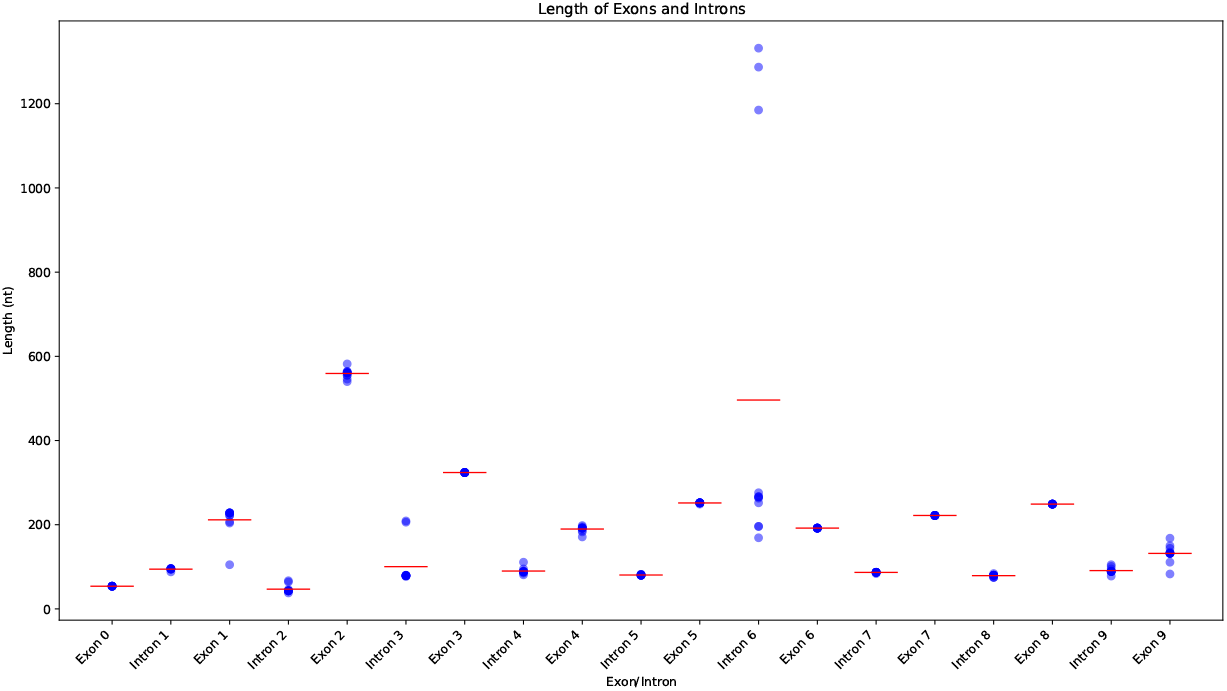
Distribution of exon and intron lengths for TERT genes across sequenced *Andrenae* species.

**Figure S4:**
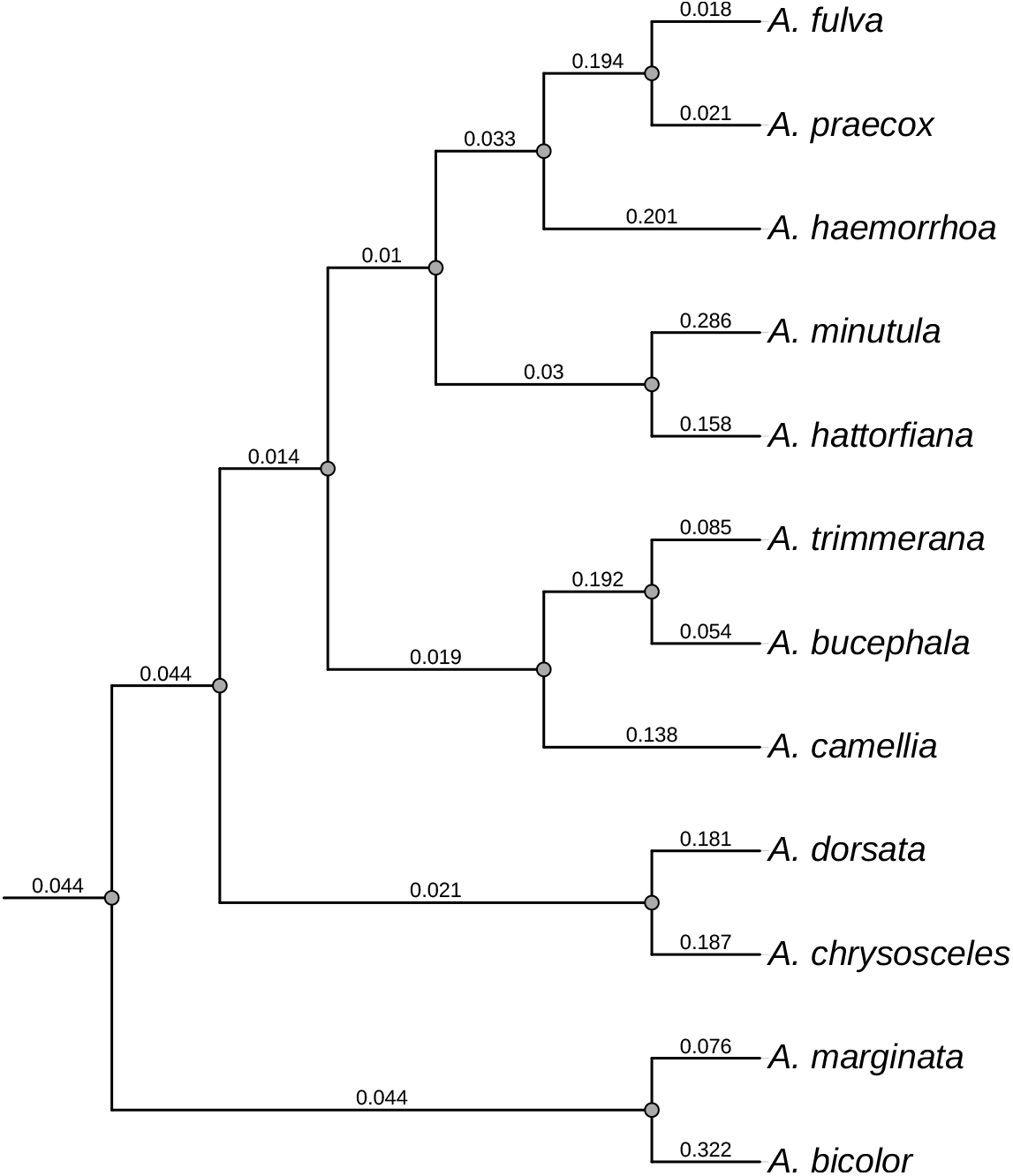
Phylogeny of *Andrena* species that we calculated based on TERT protein sequence alignments using `IQtree` Nguyen et al. [2015]. It serves as a comparison to the accepted phylogeny and was used to place *A. praecox* and *A. camellia* in the tree.

**Figure S5:**
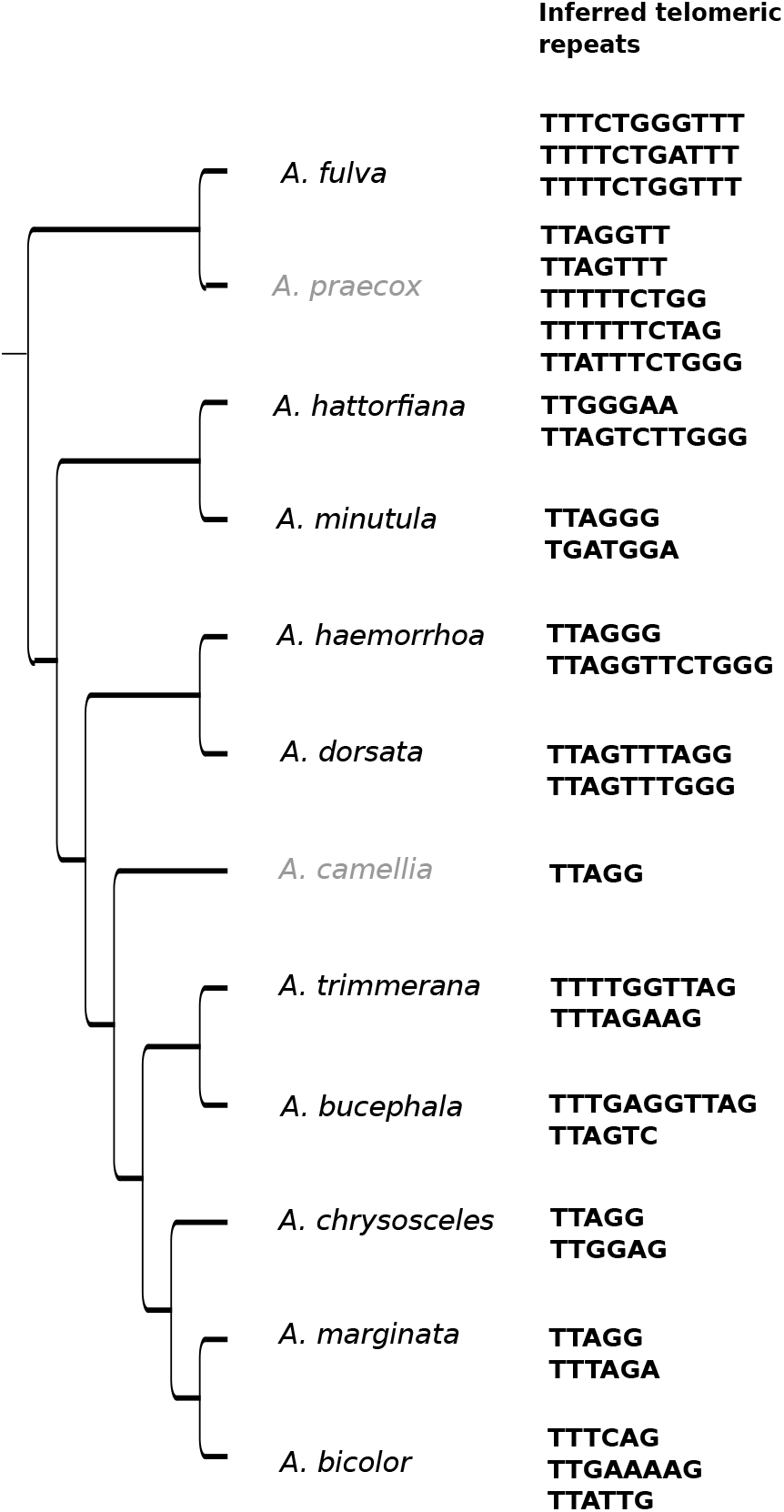
Phylogeny of *Andrena* species with *A. praecox* and *A. camellia* added. We listed predicted telomeric tandem repeats on the right.

**Figure S6:**
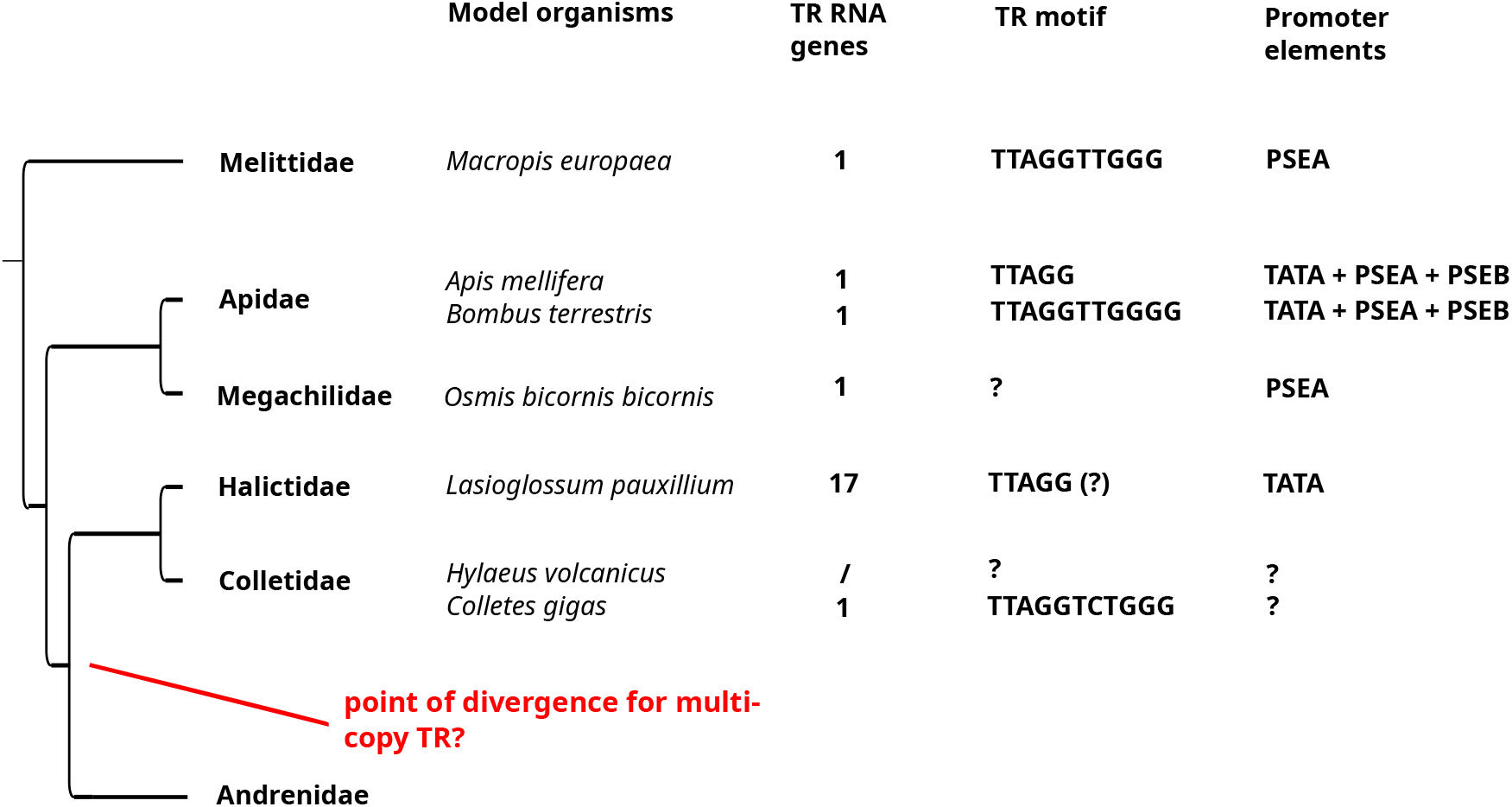
Accepted phylogeny and current understanding of TR genes and telomeric tandem repeats in closely related Hymenoptera. In none of the model organisms chosen for these more distant clades could we identify more than one TR. However, in the closely related *Lasioglossum pauxillium* we identified a large number of putative homologous TR genes, indicating the existence of TR duplicates in this organism as well. We doubt, however, that all of these are genuine and assume that some may be a product of faulty genome assembly. Furthermore, no conclusive evidence of concurrent telomeric tandem repeats was found. An analysis of promoter regions in these species (compare Figure S7) identified *Apidae* as the most similar in terms of promoter structure. The estimate of the most likely point of divergence of multi-copy TR in the tree is indicated. This is derived from the observation of *Apidae, Megachilidae* and *Melittidae* being single-copy, while we observe multiple TR loci in *Lasioglossum pauxillium*.

**Figure S7:**
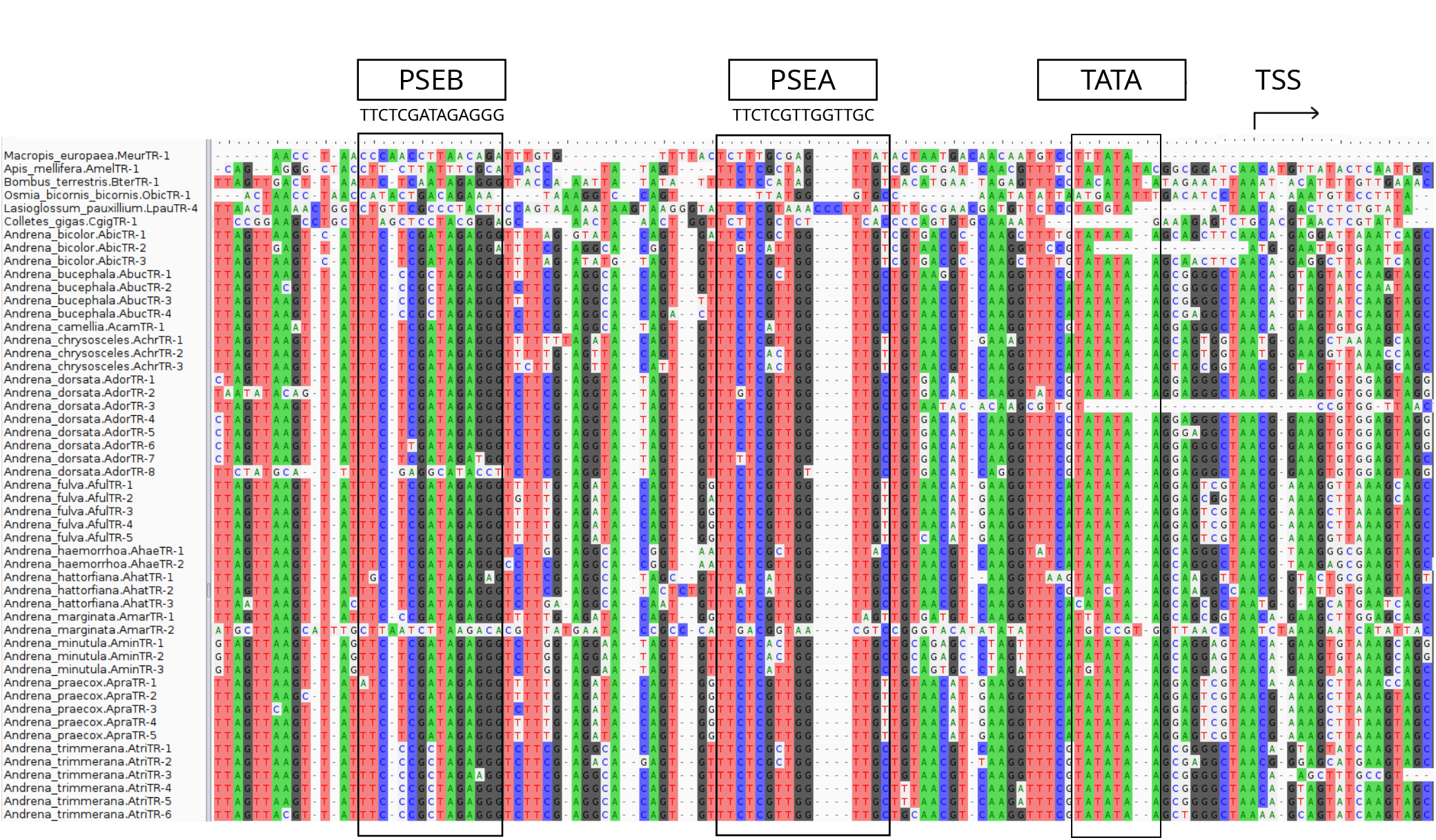
Alignment of TR candidate promoters relative to the predicted transcription start site (TSS). A conserved element 35 nt upstream of the TATA box that closely matches a PSEA promoter element starting with TTCTC as characterized by Mount et al. [2007] in *Apis mellifera* is observable. There is another conserved motif upstream of *Andrena* TR genes, hinting at the presence of a promoter proximal sequence element B (PSEB). Notable difference in promoter structure appear in the upstream regions of *Macropis europaea, Osmia bicornis bicornis* and *Colletes gigas*, all TR genes previously annotated by Fajkus et al. [2023]. Order of species in the alignment is either according to the phylogeny depicted in Figure S6 or alphabetically for *Andrena*.

**Figure S8:**
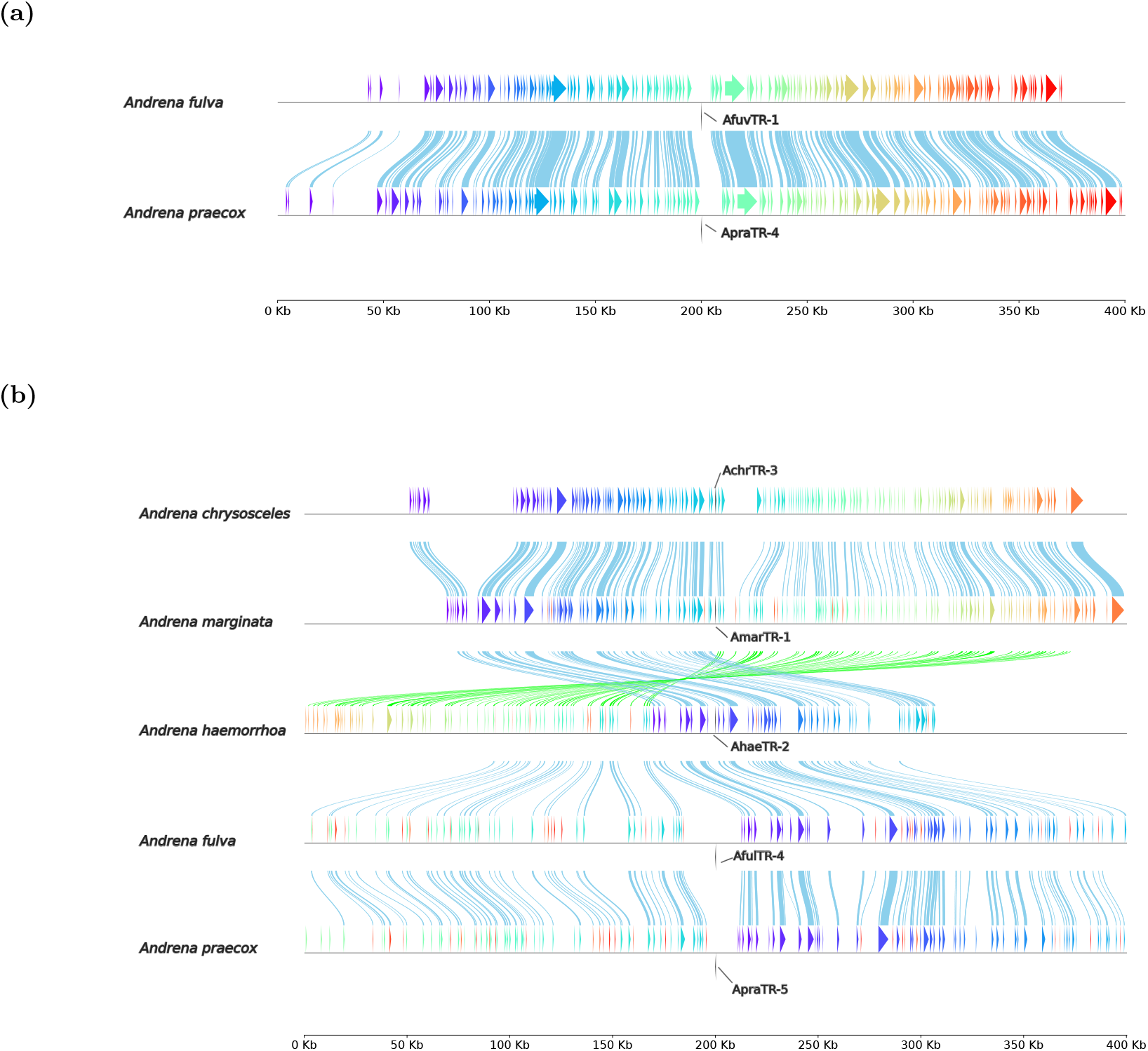
Illustration of syntenic clusters II and VI within a range of 400 kb. Directly aligned synteny anchors are highlighted and connected for clarity. The numbering of TR genes refers to their position in Table 1 for this species. (a) Cluster II contains TR genes of *A. fulva* and *A. praecox*. (b) Cluster VI serves as an interesting illustration of genomic rearrangement happening at some point during history between *A. marginata* and *A. haemorrhoa*. Judging by the new positioning of genetic loci around the TR gene it appears that *A. chrysosceles* and *A. marginata* are more closely related with each other than with the other three species shown here, which in turn show more similarity among themselves. The coexistence of these two clusters in the same species, i.e. *A. fulva* and *A. praecox*, supports the assumption that the respective TR genes being independent evolutionary products.

**Figure S9:**
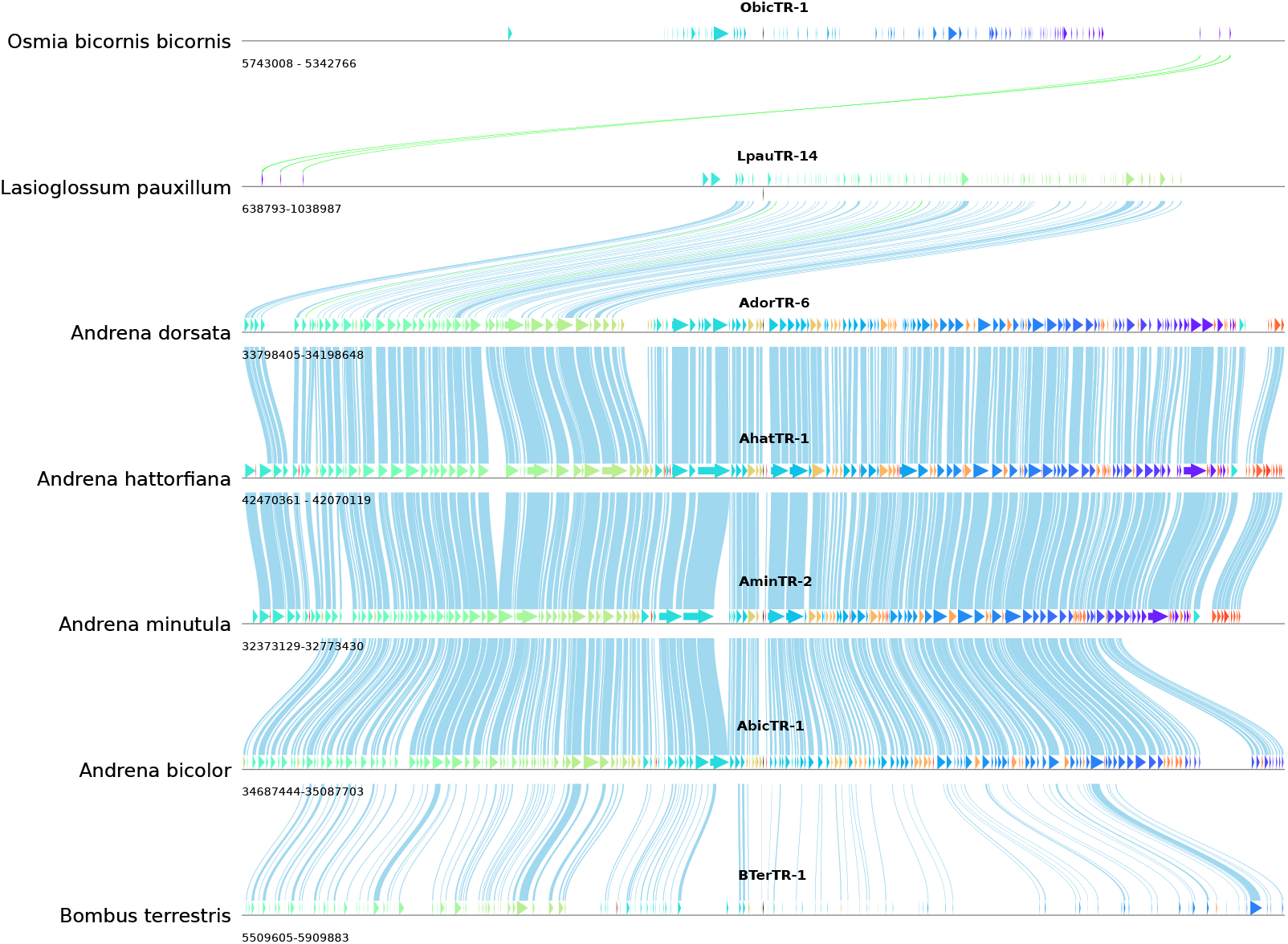
Expanded syntenic cluster analysis including TRs in related Hymenoptera. We note a conservation of genomic environment in *Bombus terrestris* and some isolated conserved regions in *Lasioglossum pauxillium* and *Osmia bicornis bicornis*.

**Figure S10:**
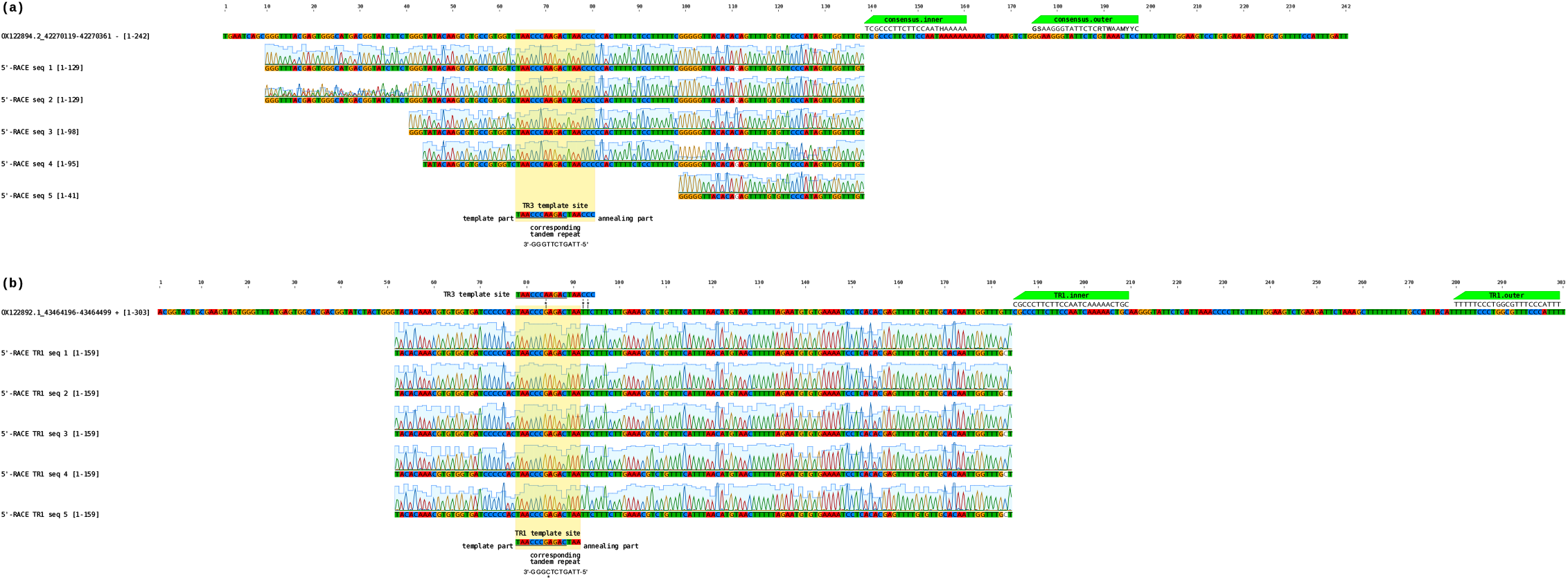
Multiple sequence alignments illustrating the expression of *A. hattorfiana* TR genes. Sequence alignments of two TR candidate genes from *A. hattorfiana* and the supporting 5’-RACE sequences, with chromatograms, each aligned relative to their putative template sites. (a) Alignment of RACE sequences (seq1 to seq5) to the candidate gene AhatTR-3 (OX122892.1, positions 43464,196–43,464,499), previously identified by Fajkus et al. [2023]. The DNA subsequence likely corresponding to the template site is highlighted in yellow (<u>TAACCCAAGAC</u>TAACCC), with the template region underlined. The resulting telomeric repeat is indicated below. (b) Alignment of RACE sequences (seq1 to seq5) to the candidate gene AhatTR-1 (OX122894.2, positions 42,270,119–42,270,361), which is presumed to be non-functional. The putative template region (<u>TAACCCGAGAC</u>TAA) of AhatTR-1 contains three mismatches compared to the AhatTR-3 template site (shown above for reference, with mismatches marked by asterisks). The telomeric tandem repeat specified by AhatTR-1 is rarely observed in DNA sequencing data, whereas the repeat specified by AhatTR-3 is among the ten most frequent tandem repeats. RACE primer positions are indicated by green boxes, with their reverse complementary sequences shown. The inability to recover the full 5’ ends of the predicted TR genes by RACE may be due to the presence of repeated triple-G regions, which serve as templates for the RACE adaptor during the initial synthesis reaction, potentially resulting in 5’-truncated RACE templates.

## Footnotes

1 https://github.com/chrisBioInf/TR-RNA-gene-duplication-supplementary-scripts

2 https://github.com/zdenkas/tandem-repeats-merger

3 https://github.com/chrisBioInf/TR-RNA-gene-duplication-supplementary-scripts

4 https://github.com/frarei312/ExceS-A-An-Exon-Centric-Split-Aligner

## References

C. Autexier and N. F. Lue. The structure and function of telomerase reverse transcriptase. Annu. Rev. Biochem., 75:493–517, 2006.

T. L. Bailey, M. Boden, F. A. Buske, M. Frith, C. E. Grant, L. Clementi, J. Ren, W. W. Li, and W. S. Noble. MEME SUITE: tools for motif discovery and searching. Nucleic acids research, 37(suppl 2):W202–W208, 2009.

T. L. Bailey, J. Johnson, C. E. Grant, and W. S. Noble. The MEME suite. Nucleic acids research, 43(W1):W39–W49, 2015.

G. Benson. Tandem repeats finder: a program to analyze DNA sequences. Nucleic acids research, 27(2):573–580, 1999.

H. Biessmann and J. Mason. Telomerase-independent mechanisms of telomere elongation. Cellular and Molecular Life Sciences CMLS, 60:2325–2333, 2003.

S. Bossert, T. J. Wood, S. Patiny, D. Michez, E. A. Almeida, R. L. Minckley, L. Packer, J. L. Neff, R. S. Copeland, J. Straka, et al. Phylogeny, biogeography and diversification of the mining bee family Andrenidae. Systematic Entomology, 47(2):283–302, 2022.

T. M. Bryan, K. J. Goodrich, and T. R. Cech. Telomerase RNA bound by protein motifs specific to telomerase reverse transcriptase. Molecular cell, 6(2):493–499, 2000.

F. Červenák, R. Sepšiová, J. Nosek, and L. Tomáška. Step-by-step evolution of telomeres: lessons from yeasts. Genome biology and evolution, 13(2):evaa268, 2021.

Y. S. Chou, D. Logeswaran, C. N. Chow, P. L. Dunn, J. D. Podlevsky, T. Liu, K. Akhter, and J. J. Chen. A degenerate telomerase RNA directs telomeric DNA synthesis in lepidopteran insects. Proc Natl Acad Sci U S A, 122(9):e2424443122, 2025. doi: 10.1073/pnas.2424443122.

M. Cohn, M. J. McEachern, and E. H. Blackburn. Telomeric sequence diversity within the genus Saccharomyces. Current genetics, 33:83–91, 1998.

S. R. Eddy. Accelerated profile HMM searches. PLoS computational biology, 7(10):e1002195, 2011.

P. Fajkus, A. Kilar, A. D. Nelson, M. Holá, V. Peška, I. Goffová, M. Fojtová, D. Zachová, J. Fulnečková, and J. Fajkus. Evolution of plant telomerase RNAs: farther to the past, deeper to the roots. Nucleic acids research, 49(13):7680–7694, 2021.

P. Fajkus, M. Adamik, A. D. Nelson, A. M. Kilar, M. Franek, M. Bubeník, R. Č. Frydrychová, A. Votavová, E. Sy‘korová, J. Fajkus, et al. Telomerase RNA in hymenoptera (insecta) switched to plant/ciliate-like biogenesis. Nucleic Acids Research, 51(1):420–433, 2023.

G. Hernandez Jr, F. Valafar, and W. E. Stumph. Insect small nuclear RNA gene promoters evolve rapidly yet retain conserved features involved in determining promoter activity and RNA polymerase specificity. Nucleic acids research, 35(1):21–34, 2007.

J. Kanoh and F. Ishikawa. Composition and conservation of the telomeric complex. Cellular and Molecular Life Sciences CMLS, 60:2295–2302, 2003.

K. Käther, S. Lemke, and P. F. Stadler. Annotation-free identification of potential synteny anchors. In International Work-Conference on Bioinformatics and Biomedical Engineering, pages 217–230. Springer, 2023.

W. J. Kent. BLAT–the BLAST-like alignment tool. Genome research, 12(4):656–664, 2002.

P. V. Kuprys, S. M. Davis, T. M. Hauer, M. Meltser, Y. Tzfati, and K. E. Kirk. Identification of telomerase RNAs from filamentous fungi reveals conservation with vertebrates and yeasts. PloS one, 8(3):e58661, 2013.

V. Kuznetsova, S. Grozeva, and V. Gokhman. Telomere structure in insects: A review. Journal of Zoological Systematics and Evolutionary Research, 58(1):127–158, 2020.

I. Letunic and P. Bork. Interactive Tree Of Life (iTOL) v5: an online tool for phylogenetic tree display and annotation. Nucleic acids research, 49(W1):W293–W296, 2021.

M. Z. Levy, R. C. Allsopp, A. B. Futcher, C. W. Greider, and C. B. Harley. Telomere end-replication problem and cell aging. Journal of molecular biology, 225(4):951–960, 1992.

G. Lin, Z. Huang, B. He, K. Jiang, T. Su, and F. Zhao. Evolutionary adaptation of genes involved in galactose derivatives metabolism in oil-tea specialized Andrena species. Genes, 14(5):1117, 2023.

D. Logeswaran, Y. Li, J. D. Podlevsky, and J. J.-L. Chen. Monophyletic origin and divergent evolution of animal telomerase RNA. Mol Biol Evol, 38(1):215–228, 2021. doi: 10.1093/molbev/msaa203.

V. A. Lukhtanov and E. A. Pazhenkova. Diversity and evolution of telomeric motifs and telomere DNA organization in insects. Biological Journal of the Linnean Society, 140(4): 536–555, 2023.

M. Mason, A. Schuller, and E. Skordalakes. Telomerase structure function. Current opinion in structural biology, 21(1):92–100, 2011.

J. Meyne, R. L. Ratliff, and R. K. MoYzIs. Conservation of the human telomere sequence (TTAGGG)n among vertebrates. Proceedings of the National Academy of Sciences, 86(18): 7049–7053, 1989.

S. M. Mount, V. Gotea, C.-F. Lin, K. Hernandez, and W. Maka-lowski. Spliceosomal small nuclear RNA genes in 11 insect genomes. RNA, 13(1):5–14, 2007.

C. Musgrove, L. I. Jansson, and M. D. Stone. New perspectives on telomerase RNA structure and function. Wiley Interdisciplinary Reviews: RNA, 9(2):e1456, 2018.

L.-T. Nguyen, H. A. Schmidt, A. Von Haeseler, and B. Q. Minh. IQ-TREE: a fast and effective stochastic algorithm for estimating maximum-likelihood phylogenies. Molecular biology and evolution, 32(1):268–274, 2015.

C. I. Nugent and V. Lundblad. The telomerase reverse transcriptase: components and regulation. Genes & development, 12(8):1073–1085, 1998.

M.-L. Pardue, S. Rashkova, E. Casacuberta, P. DeBaryshe, J. George, and K. Traverse. Two retrotransposons maintain telomeres in Drosophila. Chromosome research, 13:443–453, 2005.

T. Paysan-Lafosse, M. Blum, S. Chuguransky, T. Grego, B. L. Pinto, G. A. Salazar, M. L. Bileschi, P. Bork, A. Bridge, L. Colwell, et al. InterPro in 2022. Nucleic acids research, 51 (D1):D418–D427, 2023.

V. Peska, P. Fajkus, M. Bubeník, V. Brázda, N. Bohálová, V. Dvořáček, J. Fajkus, and S. Garcia. Extraordinary diversity of telomeres, telomerase RNAs and their template regions in Saccharomycetaceae. Scientific Reports, 11(1):12784, 2021.

G. Pisanty, R. Richter, T. Martin, J. Dettman, and S. Cardinal. Molecular phylogeny, historical biogeography and revised classification of andrenine bees (Hymenoptera: Andrenidae). Molecular Phylogenetics and Evolution, 170:107151, 2022.

J. D. Podlevsky and J. J.-L. Chen. Evolutionary perspectives of telomerase RNA structure and function. RNA biology, 13(8):720–732, 2016.

F. Reinhardt and P. F. Stadler. ExceS-A: an exon-centric split aligner. Journal of integrative bioinformatics, 19(1):20210040, 2022.

J. Song, D. Logeswaran, C. Castillo-González, Y. Li, S. Bose, B. B. Aklilu, Z. Ma, A. Polkhovskiy, J. J.-L. Chen, and D. E. Shippen. The conserved structure of plant telomerase RNA provides the missing link for an evolutionary pathway from ciliates to humans. Proceedings of the National Academy of Sciences, 116(49):24542–24550, 2019.

J. Sun, X. Li, X. Hou, S. Cao, W. Cao, Y. Zhang, J. Song, M. Wang, H. Wang, X. Yan, et al. Structural basis of human SNAPc recognizing proximal sequence element of snRNA promoter. Nature Communications, 13(1):6871, 2022.

C. A. Theimer and J. Feigon. Structure and function of telomerase RNA. Current opinion in structural biology, 16(3):307–318, 2006.

M. Waldl, B. C. Thiel, R. Ochsenreiter, A. Holzenleiter, J. V. de Araujo Oliveira, M. E. M. Walter, M. T. Wolfinger, and P. F. Stadler. TERribly difficult: Searching for telomerase rnas in Saccharomycetes. Genes, 9(8):372, 2018.

D. Wynford-Thomas and D. Kipling. The end-replication problem. Nature, 389(6651):551–551, 1997.

M. Xie, A. Mosig, X. Qi, Y. Li, P. F. Stadler, and J. J.-L. Chen. Size variation and structural conservation of vertebrate telomerase RNA. J. Biol. Chem, 283:2049–2059, 2008.

M. Zavodník, P. Fajkus, M. Franek, D. Kopecky‘, S. Garcia, S. Dodsworth, A. Orejuela, A. Kilar, J. Ptáček, M. Mátl, et al. Telomerase RNA gene paralogs in plants–the usual pathway to unusual telomeres. New Phytologist, 239(6):2353–2366, 2023.

Y. Zhou, Y. Wang, X. Xiong, A. G. Appel, C. Zhang, and X. Wang. Profiles of telomeric repeats in Insecta reveal diverse forms of telomeric motifs in Hymenopterans. Life Science Alliance, 5(7), 2022.

